# Differential Analysis of Gene Spatial Organisation with Minkowski Functionals and Tensors

**DOI:** 10.64898/2026.05.12.724373

**Authors:** Philippe Baratta, Paul Villoutreix, Anaïs Baudot

## Abstract

Spatial transcriptomics measures gene expression together with transcript coordinates in tissues. To date, comparing spatial gene expression patterns within and across samples remains challenging. We present here `minkiPy`, a geometric framework that computes, for each gene, a compact profile of morphological and topological descriptors based on Minkowski functionals and tensors. These profiles are defined in a shared feature space, enabling direct comparison of spatial organisation across genes, samples, and conditions, and the ranking of genes by the magnitude of their spatial reorganisation. We applied `minkiPy` to a MERFISH dataset of control and facioscapulohumeral muscular dystrophy myoblast cultures and to a Visium HD dataset of colorectal cancer and normal adjacent tissues, illustrating its utility across tissue types and spatial transcriptomics platforms. `minkiPy` is an open-source Python library available at https://github.com/BAUDOTlab/minkiPy.

## 1 Introduction

Spatial Transcriptomics (ST) couples gene expression measurements with spatial coordinates in tissue sections [1]. Current ST platforms generally belong to two families. Imaging-based assays, such as MERFISH [2], resolve individual transcripts in situ from predefined gene panels. Sequencing-based assays, such as Visium [3], profile transcriptomes on barcoded spots with assay-dependent spot size. Together, these ST technologies span subcellular to multicellular scales and now constitute a widely adopted experimental framework [4].

The analysis of ST data typically focuses on characterising tissue architecture at different levels of organisation [4]. Domain detection methods, for instance, aim to group spatial locations into areas composed of locally homogeneous expression patterns [5]. Another set of classical analyses aims to infer local cell–cell communications, for example by quantifying ligand–receptor co-expression [6]. Finally, ST downstream analyses often consider the detection of spatially variable genes (SVGs), transcripts that exhibit statistically significant spatial variation [7].

Different strategies further aim to organise SVGs into archetypal spatial patterns [4]. Current methods generally address this either by clustering genes using spatial dependence measures, such as spatial auto- or cross-correlation, or by learning latent representations that capture joint expression and neighbourhood structure, potentially integrating histological information [8, 9, 10]. However, to the best of our knowledge, existing methods provide only limited support for directly quantifying how the spatial organisation of individual genes differs across tissues or experimental conditions.

Different strategies have been developed recently to compare spatial organisation across samples. First, spatial alignment and registration algorithms, such as `PASTE` [11] and `STalign` [12], map tissue samples onto a common coordinate system. These methods facilitate visual and structural alignment across samples, which can be used before downstream comparative analyses. However, they do not provide explicit quantifications of spatial differences and may be unreliable when matching regions cannot be identified across samples, as in comparisons between tumour and healthy tissues [13]. A second category of methods, including `PRECAST` [14] and `DR-ST` [15], has been developed to compare global spatial organisation across samples. They project ST data from multiple samples into a common latent feature space. Latent representations support integrated analyses, including feature clustering and cell-type annotation. However, these approaches do not characterise gene-level spatial patterns nor their changes across conditions. A third category of methods, such as `DESpace` [16] and `SPADE` [17], focuses on statistical testing of gene-level spatial variation. These methods define spatial structures through specific statistical representations, such as spatial clusters or covariance kernels, and perform gene-level tests within or between samples. However, they mainly return statistical evidence for spatial differences, rather than a direct measure of the magnitude of gene-level spatial reorganisation across conditions. Consequently, current workflows lack a direct metric to characterise and quantify gene spatial reorganisation across samples and conditions. Ideally, such a metric should provide a comprehensive description of spatial organisation, encompassing global morphology (extent, shape and anisotropy), topological structure (connectivity, holes and fragmentation), scale-dependent structure, and spatial correlation dependence (clustering and dispersion), while remaining interpretable and comparable across samples and conditions.

Minkowski functionals and tensors provide precisely such descriptors. Rooted in integral geometry, Minkowski functionals form a complete family of additive measures, namely area, boundary length, and Euler characteristic. Together with their tensorial extensions, which capture directional information, they provide a compact description of morphology, anisotropy, topology, and correlation structure in spatial distributions [18, 19, 20, 21, 22]. Minkowski functionals have already demonstrated utility across biomedical imaging applications, from quantifying tumour heterogeneity to characterising tissue architecture [23, 24, 25]. However, to the best of our knowledge, Minkowski functionals or tensors have not yet been applied to spatial (transcript)omics data.

Here, we propose a gene-centred geometric framework for ST based on Minkowski functionals and one Minkowski tensor. For each gene in each sample, we treat expression as a spatial field and summarise it by a Minkowski profile. This Minkowski profile yields a compact description of the spatial organisation of each gene, capturing occupied extent, boundary complexity, fragmentation, directional bias and correlation structure. Because the construction is identical for all genes and all samples, the gene Minkowski profiles lie in a shared feature space in which distance metrics can be defined, enabling comparisons between genes within a given sample as well as between genes across samples and conditions. In addition, we quantify the uncertainty associated with each gene’s Minkowski profile using Monte Carlo resampling and incorporate these estimates into the distance metrics.

We implement this framework as an open-source Python library, `minkiPy` (https://github.com/BAUDOTlab/minkiPy), which provides an end-to-end pipeline compatible with both imaging-based and sequencing-based ST technologies. We illustrate `minkiPy` with a Facioscapulohumeral muscular dystrophy MERFISH dataset [26] and a colorectal cancer Visium HD dataset [27].

## 2 Results

### 2.1 The minkiPy framework

The overall strategy of `minkiPy` is to treat the expression of each gene in each sample as a spatial field and summarise it by a Minkowski profile (Figure 1, Methods). This profile is defined as a vector formed by the values of four Minkowski characteristics (three scalar Minkowski functionals and one tensor-derived anisotropy index) evaluated across a series of density level sets. Minkowski profiles provide a rich mathematical description of each gene’s expression morphology, topology, and correlation structure. Importantly, because we apply the same Minkowski profile construction and the same level sets to all genes, all Minkowski profiles live in a common feature space, irrespective of the sample, condition, or whether the gene was measured with imaging- or sequencing-based spatial omics technologies. In this feature space, each gene is thus represented by a geometric fingerprint (the Minkowski profile), and distances between the Minkowski profiles can be computed to compare spatial patterns within and across samples. The workflow is described here, while detailed mathematical definitions, notations, and implementation choices are provided in the Methods section.

**Figure 1:**
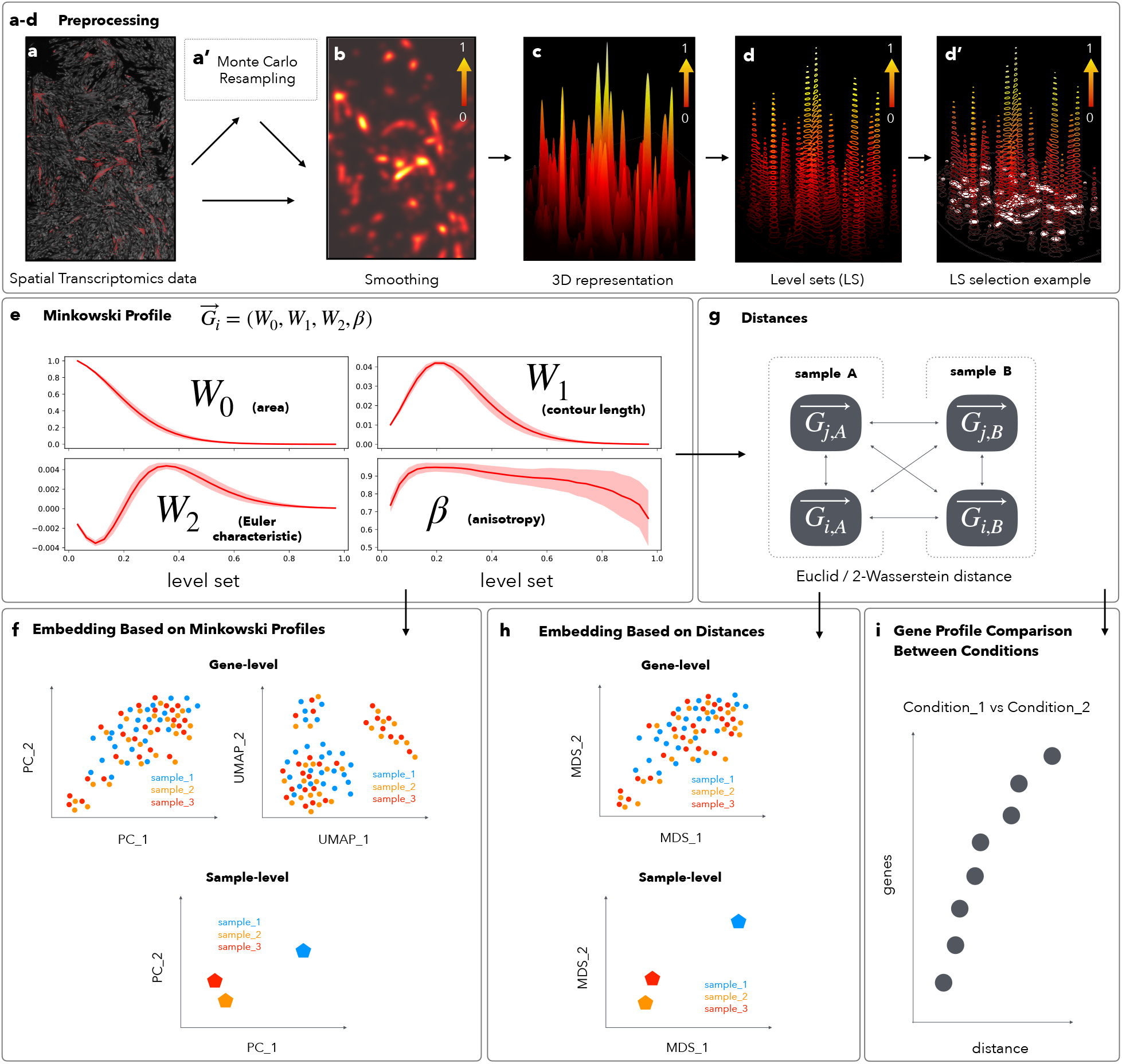
Overview of the `minkiPy` framework. A raw discrete transcript point distribution of a given gene **(a)** is transformed by Gaussian smoothing into a continuous density field, which is divided by its mean value and then normalised to the interval [0,1] **(b). (c)** displays the continuous density field of **(b)** as a three-dimensional map. In **(d)**, the field is thresholded at multiple density values, called level sets. This produces a sequence of excursion sets across the level sets. The sequence of excursion sets serves as the basis to characterise the spatial geometry of the gene across density values. **(d’)** shows an excursion set at a randomly selected level set for illustration. In **(e)**, `minkiPy` computes, for a given gene, at each level set, four descriptors of the spatial structure of the excursion set: *W*_0_ (area fraction), *W*_1_ (contour length), *W*_2_ (Euler characteristic), and *β* (tensor-derived anisotropy index). These four descriptors are referred to as the Minkowski characteristics (MCs). Optionally, `minkiPy` applies Monte Carlo resampling **(a’)** to generate multiple stochastic realisations of the initial transcript distribution, which are used to estimate the covariance matrix associated with the gene’s Minkowski profile. This covariance matrix contains uncertainties, shown in **(e)** as ±1 standard deviation. Each descriptor is then robust-scaled, and concatenating (*W*_0_, *W*_1_, *W*_2_, *β*) across all level sets yields the Minkowski profile G_*i*_ of the gene *i*, which describes its spatial structure. A Minkowski profile is computed for each gene in each sample. Profiles can be averaged, e.g., across replicate samples of the same condition. In **(f)**, embeddings are computed directly from the Minkowski profiles to explore gene and sample global spatial organisation. At the gene level, embeddings are computed using Principal component analysis (PCA) and Uniform Manifold Approximation and Projection (UMAP), and each point represents one gene in one sample. At the sample level, only PCA embedding is computed, and each point represents one sample. In **(g)**, distances between gene Minkowski profiles are computed within and across samples or averaged conditions. Here, *i* and *j* denote genes, and *A* and *B* denote samples. Distances can be Euclidean distances between robust-scaled Minkowski profiles, or 2-Wasserstein distances when Monte Carlo covariances are taken into account. In **(h)**, classical multidimensional scaling (MDS) is applied to these distance matrices at the gene or sample levels. In **(i)**, genes are ranked according to the distance between their Minkowski profiles across samples/conditions.

#### Data preprocessing

Imaging-based ST data provide individual transcript coordinates whereas sequencing-based ST data report counts on a regular grid. For sequencing-based datasets, we represent each spot by its centroid and assign its measured transcript counts to that location. Both imaging-based and sequencing-based data types are thus reduced to discrete spatial distributions of transcript locations (Figure 1a). We then transform these discrete locations into a Gaussian-smoothed density field on a regular grid, which yields a continuous representation of local transcript density (Figure 1b and c). We next divide this field by its spatial mean to obtain a contrast field that reduces the influence of overall expression level on the description of gene spatial organisation. This contrast field is then normalised to the interval [0, 1] so that all genes share a common intensity range and can be analysed with the same set of density thresholds (called level sets in our framework). We threshold the normalised field at multiple level sets, and at each level set, the field is partitioned into “in” and “out” regions that form the excursion set (Figure 1d and d’). Over all the level sets, we obtain a sequence of excursion sets (see Methods section 4.1 for details).

#### Minkowski functionals and tensor computation

At each level set, we compute four Minkowski characteristics (three scalar Minkowski functionals and one tensor-derived anisotropy index, Figure 1e). *W*_0_ measures the area occupied by the excursion set and thus captures the spatial extent of gene expression. *W*_1_ quantifies boundary length and probes the intricacy and fragmentation of gene expression. *W*_2_ is proportional to the Euler characteristic and reflects the topological organisation of expression domains, distinguishing patterns made of isolated clusters, percolating networks, or perforated structures. Beyond their geometric interpretation, these three functionals jointly contain information about the correlation structure of the distribution. The anisotropy index *β* summarises directional information encoded in the Minkowski tensor 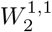, with values close to 1 indicating isotropic patterns and decreasing values reflecting increasing anisotropy. Importantly, this strategy makes no assumption about the type of spatial gene expression distribution.

#### Minkowski profile construction

For each gene, we concatenate the values of G = (*W*_0_, *W*_1_, *W*_2_, *β*) across all level sets into a Minkowski profile. The Minkowski functionals and the tensor-derived anisotropy index have distinct physical units and span different orders of magnitude. To make them comparable and prevent any single descriptor from dominating downstream analyses, we apply global robust scaling across genes and samples. Each characteristic is centred by its median and scaled by its interquartile range. This yields a robust-scaled Minkowski profile vector space, i.e. a feature space shared by all genes. This feature space can be used for gene-based embeddings using PCA or UMAP and also to compute distances across genes and samples. This common feature space also allows Minkowski profiles to be averaged, e.g., across replicate samples of the same condition.

#### Gene distance computation

As mentioned above, we use the feature space of gene Minkowski profiles to compute distances between genes within and across samples and conditions (Figure 1g). More precisely, these distances can be computed between different genes within a sample or within an averaged condition, between the same gene across samples/conditions, or between different genes across samples/conditions. These distances can then be used to cluster and compare gene patterns. We can quantify spatial pattern similarity between genes using the Euclidean distance between their robust-scaled Minkowski profiles.

To quantify uncertainty, `minkiPy` can additionally estimate, for each gene, the covariance matrix of its Minkowski profile using Monte Carlo resampling (Figure 1a’). This covariance matrix captures uncertainties and correlations between Minkowski characteristics and level sets. When the covariance matrix is considered, distances between genes are computed using a 2-Wasserstein distance that compares both profile means and their associated uncertainties and correlations. We adopt this 2-Wasserstein distance throughout the remainder of this manuscript.

### 2.2 Application of minkiPy to the Facioscapulohumeral Muscular Dystrophy Imaging-based MERFISH Dataset

In this section, we apply `minkiPy` to a publicly available ST dataset investigating facioscapulohumeral muscular dystrophy type 1 (FSHD1), a muscular dystrophy characterised by progressive weakening of facial, shoulder girdle, upper arm, and lower extremity muscles [26]. The prevailing model of FSHD1 pathogenesis implicates the aberrant expression of the transcription factor DUX4, driven by contractions of the D4Z4 repeat array on chromosome 4q35.

Chen et al. [26] used multiplexed error-robust fluorescence *in situ* hybridization (MERFISH) to spatially profile the transcriptomes of myoblast cultures across conditions. We analyse ten samples spanning three conditions: four control samples derived from myoblast cells carrying a normal number of D4Z4 repeats (∼ 20), four samples from patient-derived FSHD1 myoblast cells characterised by a contracted D4Z4 region (∼ 2 repeats), and two samples from isogenic D4Z4-contraction mutants (∼ 1 repeat) derived from the control myoblast cells, referred to as DEL5. The MERFISH panel comprises 140 genes. We apply `minkiPy` to this MERFISH dataset.

We first compute the Minkowski profile of each gene in each sample, capturing its spatial organisation at the individual sample level. We also compute an averaged Minkowski profile for each gene per condition. These averaged profiles are obtained by averaging the Minkowski profiles of a given gene across all replicate samples of a given condition. We treat the resulting averaged profiles as an additional sample. We quantify how the organisation of a given gene varies across samples or averaged conditions by computing the 2-Wasserstein distance between its Minkowski profiles (see Methods, Section 4.7). To obtain sample–sample distances, we compute the gene 2-Wasserstein distances between two samples. We combine the obtained distances in a single distance value for each pair of samples (see Methods, Section 4.9). We visualised this distance matrix using classical multidimensional scaling (MDS), which places samples close together when their overall spatial gene organisation is similar. The resulting embedding displays a structure consistent with the experimental design (Figure 2a): Control, FSHD1, and DEL5 samples segregate into distinct regions of the embedding space and the averaged samples are positioned near the centres of their respective sample groups. Similar results are obtained with a sample-level PCA embedding based on the gene Minkowski profiles (Supplementary Figure S1a).

**Figure 2:**
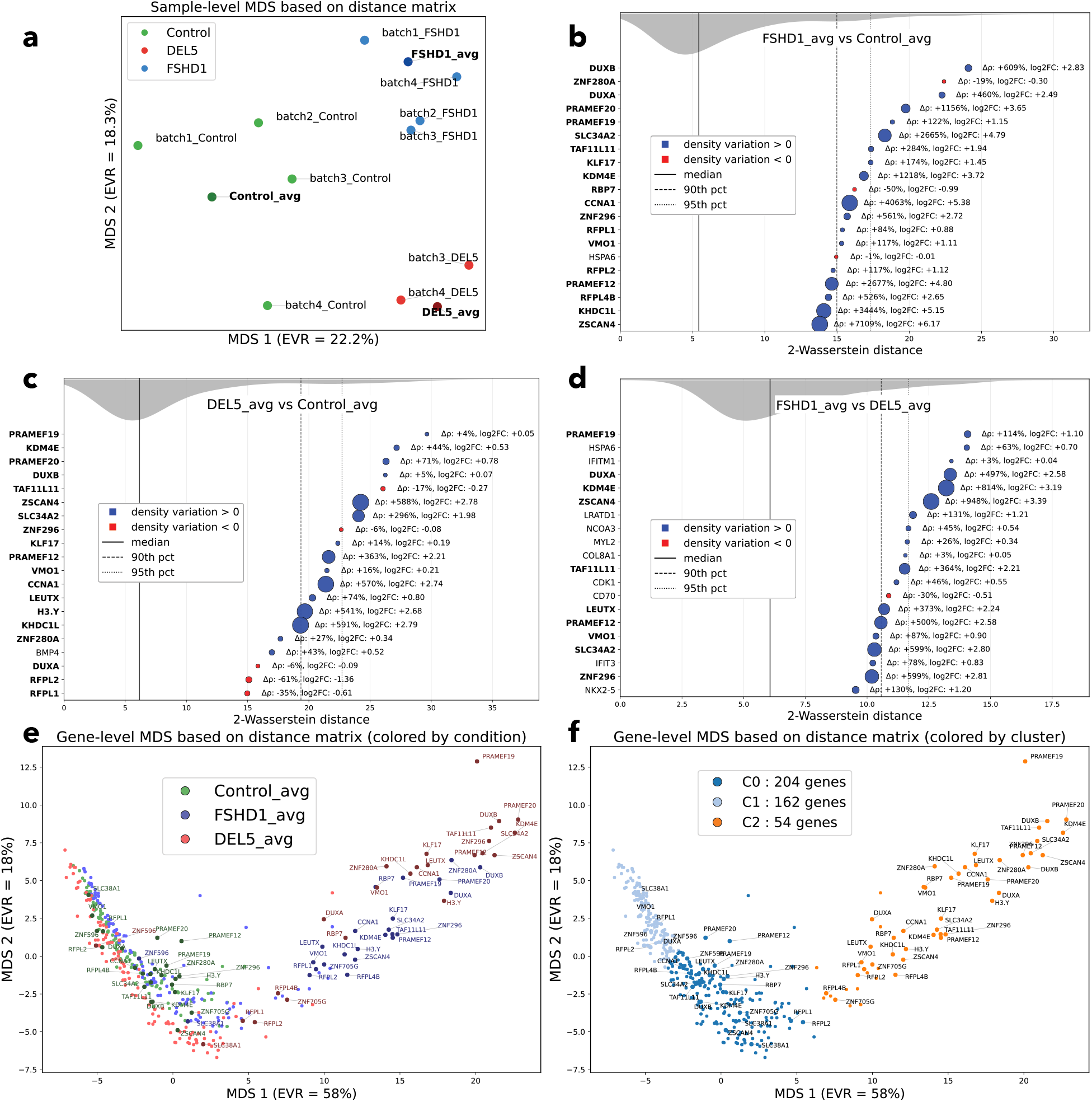
Application of `minkiPy` to the FSHD MERFISH dataset. **(a)** Sample-level classical multidimensional scaling (MDS) of the 2-Wasserstein distances between the Minkowski profiles of each gene in every pair of samples. Samples are coloured by condition, with darker points indicating the averaged samples for each condition. **(b–d)** Top 20 genes ranked by their 2-Wasserstein distances between averaged conditions. For each gene, marker position encodes its 2-Wasserstein distance between the compared averaged conditions, marker colour indicates the sign of the log_2_ fold change/normalised density (blue: higher normalised density in the first condition than in the second; red: higher normalised density in the second than in the first), and marker thickness scales with the absolute percent density change. DUX4-target genes are highlighted in bold. The grey band on the top shows the smoothed distribution of same-gene 2-Wasserstein distances across all genes, with vertical lines marking the median, 90^th^ and 95^th^ percentiles. **(e)** Classical MDS embedding of gene-level 2-Wasserstein distances, coloured by condition. Each point represents one gene in one averaged condition. Each gene therefore appears three times, once per averaged condition. EVR, explained variance ratio, is shown on the axes. DUX4-target genes are shown with darker markers and labelled. **(f)** Classical MDS of the same distance matrix, coloured by communities identified with the Leiden algorithm (resolution 0.05) on a *k*-nearest-neighbour graph with *k* = 25.

As a control, we assessed whether the observed sample segregation is also identified using only gene expression densities without considering information about spatial organisation. A PCA using, for each sample, the vector of normalised gene densities does not reveal a structure consistent with the experimental design (Supplementary Figure S1b). This indicates that the signal captured by `minkiPy` with the Minkowski-based spatial representations is not solely explained by global changes in gene expression but arises from condition-specific alterations in spatial gene organisation.

We next ask which genes display the strongest spatial differences between conditions. To this end, we use the 2-Wasserstein distances computed between each gene’s averaged Minkowski profiles in the FSHD1 versus Control, DEL5 versus Control, and FSHD1 versus DEL5 conditions (Figure 2b, c, d and Supplementary Data 1). Importantly, many of the genes top-ranked by `minkiPy` based on the distances between their Minkowski profiles also show increased normalised densities or higher log_2_ fold change across conditions (Figure 2b–d). However, correlation is weak to moderate between the gene 2-Wasserstein distances and log_2_ fold change (see Methods, Section 4.10 and Supplementary Figures S2a, b and c). This provides additional evidence that the 2-Wasserstein geometry derived from the Minkowski profiles captures aspects of spatial gene organisation that are not reducible to gene expression changes alone, and therefore provides complementary information.

Among the 140 genes in the panel, 24 genes are annotated as DUX4-target genes [26]. The DUX4 transcription factor is lowly expressed and often undetectable in tissues, and DUX4-target genes (identified by transcriptomics analyses) are considered as biomarkers of DUX4 activity. In the FSHD1 versus Control and DEL5 versus Control comparisons, we observe that the top 20 genes ranked by `minkiPy` as showing the strongest spatial reorganisation are predominantly DUX4-target genes (Figures 2b and c, DUX4-target genes in bold). This suggests that DUX4-targets contribute to FSHD1 pathophysiology not only through altered expression but also through spatial reorganisation. In addition, the top-ranked genes in the FSHD1 versus Control and DEL5 versus Control comparisons show substantial overlap, indicating that the DEL5 model recapitulates key spatial changes observed in the disease. Overlapping top-ranked genes include, for instance, *DUXA* and *DUXB*, which both show differential spatial organisation in the FSHD1 versus Control and DEL5 versus Control comparisons. *DUXA* and *DUXB* encode DNA binding proteins. *DUXA* is directly activated by DUX4 and encodes a competitive inhibitor. Interestingly, this differential spatial organisation is associated with an increased normalised expression density in FSHD1 versus Control but not in DEL5 versus Control (Supplementary Figures S3 and S4 for DAPI–transcript overlays across samples). *ZNF280A* is also an interesting gene, showing one of the strongest spatial reorganisations according to its 2-Wasserstein distance in both the FSHD1-versus-Control and DEL5-versus-Control comparisons, despite modest and opposite changes in expression density. *ZNF280A* encodes a zinc-finger protein that may act as a transcription factor (Supplementary Figure S5). A similar pattern is observed for *TAF11L11*, encoding a TATA-box binding protein with poorly described functions (Supplementary Figure S6). Finally, another interesting example is *PRAMEF19*. It shows spatial reorganisation in both the FSHD1 versus Control and DEL5 versus Control comparisons, but also in the FSHD1 versus DEL5 comparison (Figure 2d). *PRAMEF19* is predicted to be involved in proteasome-mediated ubiquitin-dependent protein catabolic process (Supplementary Figure S7).

As stated above, the genes top-ranked by `minkiPy` in the FSHD1 versus Control and DEL5 versus Control comparisons are similar, demonstrating the relevance of the DEL5 mutant model. However, we can pinpoint some differences between the disease and its model thanks to the direct comparison of FSHD1 versus DEL5 averaged conditions (Figure 2d). For instance, *HSPA6*, encoding a heat shock protein involved in stress response, shows different spatial organisation between the FSHD1 and DEL5 conditions (Supplementary Figure S8).

To obtain a global view of gene spatial organisation changes across conditions, we compute distances between all genes’ averaged Minkowski profiles (see Methods, Section 4.7). In a classical MDS embedding of the obtained gene distance matrix (see Methods, Section 4.9 and Figure 2e), two points that lie close to each other correspond to genes with similar spatial patterns whereas distant points correspond to genes with different spatial patterns. Clustering analysis reveals three communities (see Methods, Section 4.8 and Figure 2f). The left part of the embedding space contains genes from all three averaged conditions (Figure 2e; Supplementary Table S1). These genes belong to the communities C0 and C1 (Figure 2f and Supplementary Data 2). Interestingly, the genes in these communities are the non-DUX4 target genes from all three conditions, together with DUX4-target genes only from the control condition. By contrast, the right part of the embedding space, corresponding to the community C2, is almost exclusively populated by genes from the FSHD1 and DEL5 conditions that are also DUX4-target genes. `minkiPy` reveals that these genes adopt a spatial pattern that is specific to the FSHD1/DEL5 states and not observed in con_eorga. Similar results are observed with PCA and UMAP embeddings of the condition-averaged gene Minkowski profiles (Supplementary Figures S1, panels c–f), as well as when studying individual samples rather than condition-averaged profiles (Supplementary Figure S9). Note that near-indistinguishable embeddings obtained with MDS and PCA are expected when the distance matrix can be approximated by a Euclidean distance matrix; see Supplementary Section S2. Overall, the analysis of the FSHD1 dataset with `minkiPy` highlights the spatial _eorganization of DUX4-target genes in FSHD1 and in the DEL5 model beyond differential gene expression alone.

### 2.3 Application of minkiPy to the Colorectal Cancer Sequencing-based Visium HD Dataset

In this section, we apply `minkiPy` to a publicly available sequencing-based ST dataset recently published by Oliveira et al. [27]. Generated using the Visium HD platform, these data profile colorectal cancer tissues (CRC) and matched normal adjacent tissues. Unlike the MERFISH FSHD dataset (Section 2.2), where we compared gene spatial patterns within a homogeneous tissue type, these Visium HD samples are heterogeneous, spanning distinct anatomical regions, from normal mucosa to invasive carcinoma. We therefore illustrate the use of `minkiPy` to quantify gene spatial reorganisation between tumour and normal tissue.

We analyse five Visium HD sections from the colorectal cancer cohort of Oliveira et al. [27]. All samples correspond to formalin-fixed, paraffin-embedded (FFPE) tissue sections profiled with whole-transcriptome Visium HD. On the array, capture probes are arranged on a continuous grid of 2 `×` 2 *µ*m barcodes without gaps, so that the tissue is effectively tiled at (sub-)cellular resolution. The five tissue sections considered consist of three colorectal carcinoma samples (P1_CRC, P2_CRC and P5_CRC) and two normal adjacent tissues (P3_NAT and P5_NAT). These sections come from four distinct patients and span different anatomical regions of the large intestine (rectum for P1_CRC, sigmoid for P2_CRC, transverse colon for P3_NAT and not specified for P5_CRC and P5_NAT) [27].

We first compute the 2-Wasserstein distances between the Minkowski profiles of each gene in every pair of samples (see Methods, Section 4.7). We obtain a sample-sample distance matrix that aggregates the 2-Wasserstein distances computed for each gene between every pair of samples (see Methods, Section 4.9). The classical multidimensional scaling (MDS) embedding of this sample-sample distance matrix does not display a structure consistent with patient identity and disease status (Figure 3a). Indeed, CRC and NAT samples do not separate in the embedding space, with samples from different disease statuses, anatomical sites, and patients broadly scattered without discernible structure. A similar absence of structure is observed with sample-level PCA based on gene Minkowski profiles (Supplementary Figure S10a). As a control, we perform PCA using per-sample vectors of normalised gene densities, which likewise reveal no evident anatomical site, patient, or disease status structure (Supplementary Figure S10b). Together, these results indicate that samples are highly heterogeneous at both the expression and spatial organisation levels. Given this heterogeneity, we did not compute condition-averaged Minkowski profiles. We focus on the matched pair P5_CRC/P5_NAT, derived from the same individual, allowing differences in Minkowski profiles to be interpreted primarily in terms of tumour versus normal adjacent tissue rather than patient or anatomical location effects.

**Figure 3:**
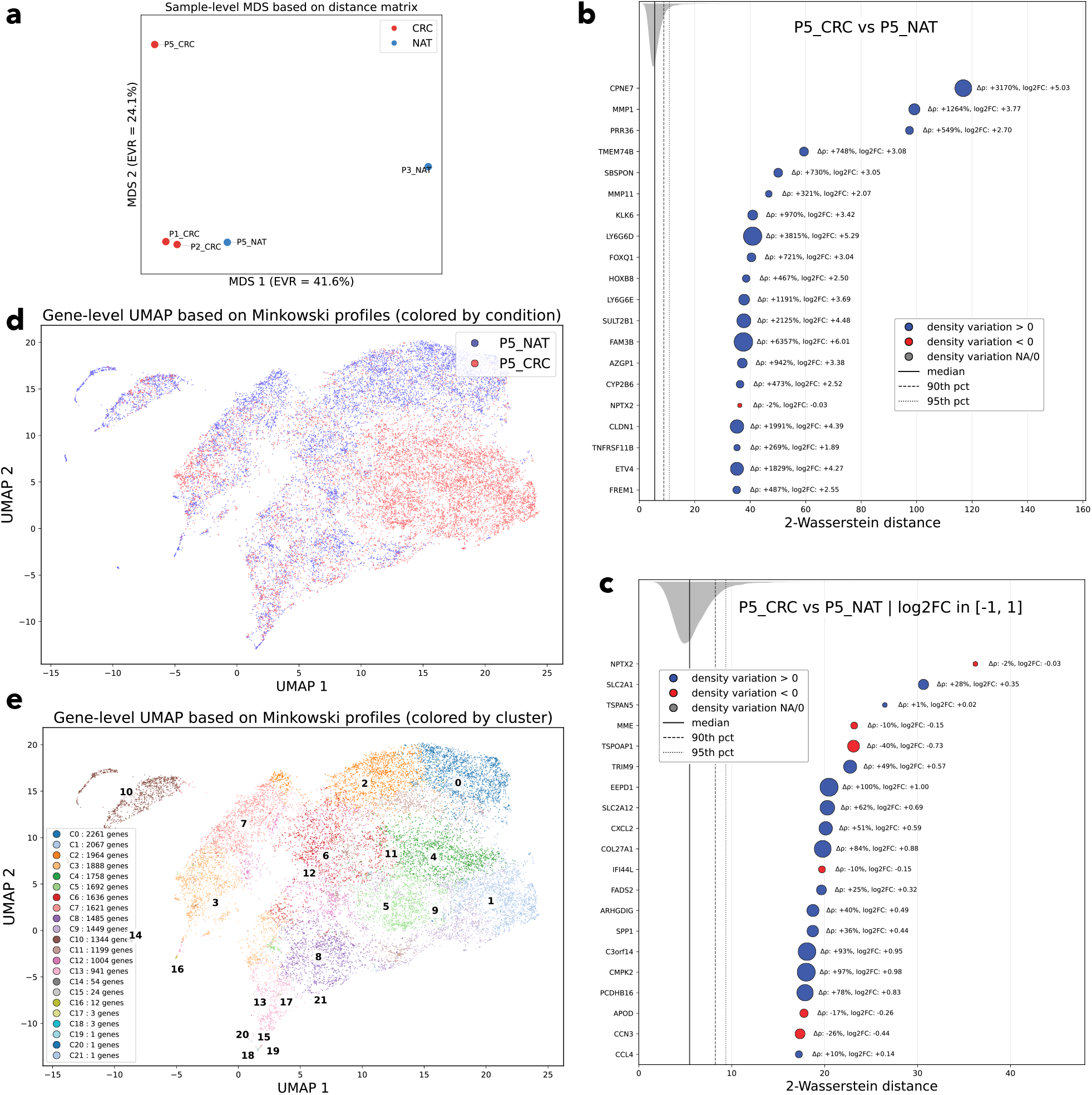
Application of `minkiPy` to the CRC Visium HD dataset. **(a)** Sample-level classical multidimensional scaling (MDS) of the 2-Wasserstein distances between the Minkowski profiles of each gene in every pair of samples. Samples are coloured by condition. **(b)** Top 20 genes ranked by their 2-Wasserstein distances between conditions (P5_CRC vs P5_NAT). For each gene, marker position encodes its 2-Wasserstein distance between the two compared samples, marker colour indicates the sign of the log_2_ fold change/normalised density (blue: higher normalised density in the first condition than in the second; red: higher normalised density in the second than in the first), and marker thickness scales with the absolute percent density change. The grey band on the top shows the smoothed distribution of same-gene 2-Wasserstein distances across all genes, with vertical lines marking the median, 90^th^ and 95^th^ percentiles. **(c)** Same representation as in **(b)** after filtering genes with log_2_ fold change between −1 and 1, highlighting genes that exhibit spatial reorganisation without substantial changes in normalised expression density between the compared conditions. **(d)** UMAP embedding of gene Minkowski profiles, coloured by condition. Each point represents one gene in one sample. Each gene therefore appears twice, once in P5_CRC and once in P5_NAT. **(e)** The same UMAP embedding, coloured by communities identified with the Leiden algorithm (resolution 0.04) on a *k*-nearest-neighbour graph with *k* = 200.

We identify genes displaying the strongest spatial differences between P5_CRC and P5_NAT, using the 2-Wasserstein distances computed between each gene’s Minkowski profiles in the two samples (P5_CRC versus P5_NAT). Given the high heterogeneity of the tissue sections studied here, the top-ranked genes based on distances between their Minkowski profiles are mostly expressed in the tumour but absent from the normal adjacent tissue, thereby combining strong fold changes in expression with distinct spatial patterns (Figure 3b and Supplementary Data 3). For instance, *CPNE7* and *LY6G6D* are specifically expressed in the tumour sample and localised in the tumour subregion. *CPNE7* encodes a calcium-dependent phospholipid-binding protein known to be overexpressed in CRC [28] (Supplementary Figure S11). Similarly, *LY6G6D* is associated with low expression and a poorly characterised function in normal tissue homeostasis but has recently attracted attention because of its overexpression in immunologically “cold” CRC [29] (Supplementary Figure S12). Other examples of genes overexpressed in the tumour sample but not restricted to the tumour subregion include *MMP1*, an interstitial collagenase that drives extracellular matrix degradation and tumour invasion [30, 31] and is expressed in small patches in the tumour and at the stromal-tumour interface (Supplementary Figure S13). *MMP11* (stromelysin-3), a less canonical matrix metalloproteinase with a role in colorectal cancer progression [30, 31], is expressed in patches in the tumour subregion (Supplementary Figure S14).

To isolate genes whose spatial organisation differs between samples independently of strong differential expression, we restricted the analysis to genes with a log_2_ fold change between −1 and 1 and ranked them by their 2-Wasserstein distances (Figure 3c and Supplementary Data 4). This filtering reveals genes with spatially distinct organisation between the two samples that are not driven by bulk expression differences. For instance, *NPTX2*, a secreted neuronal gene identified as associated with colorectal cancer progression [32], shows low, scattered expression throughout the normal mucosa but higher, localised expression at the tumour margin (Supplementary Figure S15). Similarly, *IFI44L*, whose expression is induced by type I interferon [33], displays low scattered expression in normal tissue but a concentrated patch within a restricted tumour subregion (Supplementary Figure S16). *SPP1*, a well-established marker of a macrophage subpopulation described in Oliveira et al. [27], provides a particularly compelling example: while it shows only sparse expression in small patches in the NAT sample, it displays strong localised expression in the CRC sample outside the tumour subregion itself, consistent with a cluster of tumour-associated macrophages (Supplementary Figure S17). Finally, *APOD*, involved in lipid metabolism and transport, shifts from a homogeneous expression pattern in the normal sample to a spatially concentrated distribution at the tumour-stroma interface in the CRC sample (Supplementary Figure S18).

To obtain a global view of gene spatial organisation, we construct a UMAP embedding of all genes’ Minkowski profiles in P5_CRC and P5_NAT (see Methods, Section 4.9, Figure 3d and e, and Supplementary Figure S10c–f for MDS and PCA). Several regions are condition-enriched. In particular, a large region of the embedding space is composed of genes from the P5_CRC sample, while several smaller regions are composed of genes from the P5_NAT sample. Clustering analysis identifies 22 communities (Figure 3e; Supplementary Table S2 and Supplementary Data 5). A subset of the communities is strongly unbalanced and contains a majority of P5_CRC genes (notably communities 1, 4, 5 and 9). Conversely, several communities are unbalanced but contain a majority of P5_NAT genes (notably communities 0, 2, 7, 10 and 11), indicating gene spatial patterns that are not present in the tumour section. We also identify balanced communities (including communities 3, 6, 8, 13). However, these communities mostly do not contain the same genes from the two conditions, as quantified by a low retention rate, i.e., the percentage of genes assigned to the same community in the two conditions (Supplementary Table S2). Community 13 has the highest retention rate, suggesting a subset of genes whose spatial organisation is comparatively stable between P5_CRC and P5_NAT. Communities are not systematically associated with higher or lower mean normalised densities, supporting the idea that community separation primarily reflects differences in spatial organisation rather than abundance alone. This corroborates the absence of a strong correlation between 2-Wasserstein distances and log_2_ fold change between the same genes in different conditions (Supplementary Figure S19).

## 3 Discussion

We developed `minkiPy` to quantify gene spatial organisation in spatial transcriptomics data. The method computes a Minkowski profile for each gene in each spatial transcriptomics sample. Minkowski profiles are built from morphological and topological descriptors that provide rich information about the spatial extent, boundary structure, topology, anisotropy and correlation structure of gene spatial organisation. The genes’ Minkowski profiles belong to a shared feature space, allowing genes’ spatial patterns to be compared within a sample, across samples, and across conditions. In addition, this allows genes’ Minkowski profiles to be averaged, for instance between different biological replicates of the same condition, between adjacent slices, or between different regions of the same sample. Importantly, `minkiPy` does not require cell segmentation, spot deconvolution, or anatomical alignment. It does not assume a specific spatial distribution and is independent of the spatial transcriptomics platform.

The comparisons of genes’ Minkowski profiles support various analyses, including the community detection of genes based on their spatial profiles or the ranking of genes by the magnitude of their spatial reorganisation. Altogether, this makes `minkiPy` a useful tool to be added to spatial transcriptomics analysis pipelines. It can be applied in addition to classical analyses (e.g., detection of spatially variable genes, detection of spatial domains) or in complement to such analyses (e.g., focusing only on spatially variable genes, to compare the gene spatial organisation of two identified regions).

We observed that `minkiPy` analysis of gene spatial organisation is not independent of gene expression. Indeed, normalising each density field as we did in `minkiPy` reduces the direct influence of total gene expression, but gene expression and spatial organisation cannot be fully decoupled. Nonetheless, in the FSHD MERFISH dataset, distances between Minkowski profiles are not strongly correlated with log_2_ fold changes. We observed similar results in the Visium HD colorectal cancer dataset. However, in the Visium HD dataset, many top-ranked genes combined strong spatial changes with strong expression differences. We therefore filtered and highlighted genes with modest log_2_ fold change and strong spatial reorganisation. These examples show that `minkiPy` is most informative when interpreted together with expression-level information. The framework is therefore complementary to differential expression analysis.

This partial coupling between gene expression and gene spatial organisation is deeply linked to the notion of shot-noise [34, 19]. `minkiPy` does not currently apply a complete shot-noise correction to the estimated amplitude of the Minkowski characteristics. This means that finite transcript sampling may still affect the profiles, even after density normalisation. We used Monte Carlo resampling to estimate the uncertainty associated with shot noise rather than to remove its effects on the amplitude of the Minkowski profile. To our knowledge, no established correction is available to remove the shot noise bias.

Other limitations could be considered. First, Minkowski profiles depend on how the density field is constructed. Grid resolution, smoothing scale and the number of level sets determine which spatial structures are captured. In practice, users should report preprocessing parameters and test whether topranked genes, embeddings and communities remain stable under moderate changes in resolution, smoothing and gene filtering.

Then, the choice of a distance metric involves a trade-off between rigour and computational cost. Euclidean distances between Minkowski profiles provide a simple baseline and could be sufficient for exploratory analyses at a reasonable computing cost. In the datasets analysed in this study, Euclidean and covariance-aware 2-Wasserstein distances give similar global embeddings. However, this may not be the case for noisier datasets or when transcript counts vary strongly across genes or samples. In such cases, uncertainty in the estimated Minkowski profiles may have stronger effects, and covariance-aware 2-Wasserstein distances may then become preferable. Computing covariance-aware 2-Wasserstein distances requires Monte Carlo resampling and covariance estimation, which can become expensive for large gene panels or whole-transcriptome datasets. Approximating the 2-Wasserstein distance using the diagonal of the covariance matrices can reduce this cost. In the datasets analysed here, this diagonal approximation produced results close to those obtained with the full covariance matrix. We recommend testing all these strategies on a subset of genes and checking whether the main rankings, embeddings and communities remain stable.

Moreover, `minkiPy` treats each gene in each sample independently. Similar Minkowski profiles indicate similar morphology/topology after preprocessing. However, similar Minkowski profiles do not necessarily imply co-localisation or spatial interaction. Two genes may have similar Minkowski profiles while occupying different tissue regions. For example, two genes expressed in alternating parallel stripes could have similar morphology, anisotropy and topology, while being spatially anti-correlated. Complementary measures, such as spatial cross-correlation, cooccurrence statistics or ligand–receptor analysis, can be applied to provide further information.

In this study, we used Minkowski profiles as summary representations of gene spatial organisation, enabling gene and sample comparison based on spatial organisation. We did not systematically decompose each gene-level distance into contributions from individual Minkowski characteristics or level sets. Indeed, in principle, each component of the profile can be inspected separately to determine whether a spatial difference is mainly driven by changes in occupied area, boundary structure, topology, anisotropy or correlation. Such component-wise analyses could provide a more detailed interpretation of spatial reorganisation and represent a natural direction for future applications of `minkiPy`.

Other applications of `minkiPy` could also consider other modalities like spatial molecular probes that can be represented as point distribution (e.g., spatial proteomics and metabolomics), imaging-derived features, or intensity fields. Interestingly, `minkiPy` could also be used to jointly study different spatial modalities. Extension to three-dimensional data is natural from the mathematical point of view, but would require volumetric density estimation, three-dimensional masks, surface-based excursion sets and adapted computational strategies. As three-dimensional spatial omics datasets become more common, this represents a clear direction for future versions of `minkiPy`.

## 4 Methods

In this section, we define the four Minkowski Char-acteristics (MCs) as the combination of three scalar Minkowski functionals and one anisotropy index derived from a Minkowski tensor. Together, the MCs allow for a comprehensive description of the morphological, topological, and correlation structures of spatial distributions [35, 21, 36]. We introduce `minkiPy`, a framework based on the computation of MCs from Spatial Transcriptomics (ST) data (Figure 1).

### 4.1 Preprocessing

Spatial transcriptomics data originate from two main technologies: imaging-based assays detect individual transcripts as point clouds and assign each a precise (*x, y*) coordinate, while sequencing-based assays report counts per spot on a regular grid. Here, for sequencing-based data, we assign all transcripts measured in a spot to the spot’s centroid. As a result, data produced by both technologies are treated as discrete spatial distributions of transcript locations.

However, computing MCs requires a binary field as input. ST data must therefore be transformed before MCs can be computed. The first step is Gaussian smoothing, which converts transcript locations into a continuous spatial signal. The result is a continuous field sampled on a regular grid with a given resolution, where each pixel encodes the local transcript density as follows:

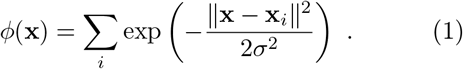

Here, *ϕ*(x) denotes the value of the continuous field at the grid-centre position x, x_*i*_ denotes the position of the *i*-th discrete data point, and *σ*^2^ is the variance of the Gaussian kernel, controlling the smoothing scale. Because both *σ* and the grid resolution jointly determine the analysis scale, we set *σ* = 2 pixels and allow the user to set the grid resolution. With *σ* fixed in pixel units, the effective smoothing scale is exclusively controlled by the grid resolution.

The grid resolution should match the intrinsic spatial resolution of the assay: it must not be finer-grained than the data themselves to avoid oversampling and discretisation artefacts, yet it must not be too coarse-grained to retain biologically meaningful spatial signal. Sensitivity analyses across a small range of resolutions can then be used to verify that the main conclusions are robust.

In addition to enabling MCs computations, Gaussian smoothing provides several advantages. First, it mitigates technology-specific errors. In imaging-based assays, these include capture inefficiency, detection sensitivity limits, and spatial misregistration or alignment artefacts from imperfectly overlaid imaging rounds or fluorescence channels, which cause slight translational or rotational shifts in the recorded positions of transcripts. In sequencing-based techniques, smoothing reduces sequencing and capture variability across spots while preserving spatial coherence across adjacent measurements [37]. Second, Gaussian smoothing diminishes sampling/shot noise in computed statistics and stabilises covariances, yielding spatial descriptors that capture true structural patterns rather than sampling artefacts [38].

Since each gene’s continuous density field *ϕ*(x) reflects both spatial organisation and overall expression level, direct comparison of Minkowski characteristics across genes would be strongly influenced by gene expression abundance differences. To reduce this influence, we first rescale the field by its spatial mean 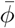, defining the contrast field:

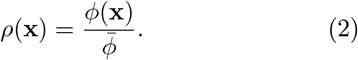

In this representation, spatial structure is described primarily by variations around the mean rather than by absolute intensity. It places density fields from different genes on a comparable scale before any further normalisation. Because the resulting contrast fields still span gene-specific ranges, we subsequently normalise each ρ(x) to the interval [0, 1]. This common scale further allows us to apply the same density thresholds when computing MCs.

By applying density thresholds, here referred to as level sets, to the continuous density fields normalised to the interval [0, 1], we generate a sequence of binary fields. Such binary fields are known as excursion sets. At a given level set, the excursion set partitions the field into “in” and “out” regions (Figure 1d’). The resulting boundaries of the excursion set are identified using the marching squares algorithm [39]. Connected “in” regions within these sets are then distinguished through a friends-of-friends algorithm [40]. The number of level sets is a user-defined parameter chosen to balance detail with computational feasibility.

### 4.2 Computing the Minkowski Functionals

Minkowski functionals provide a mathematical framework to characterise the morphology and topology of spatial fields [18, 41, 42] and capture information related to their n-point auto-correlation structure [21]. Minkowski functionals are rooted in a classical result of integral geometry known as Hadwiger’s theorem. This theorem states that in a *d*-dimensional Euclidean space, any continuous, translation- and rotation-invariant, additive valuation can be expressed as a linear combination of exactly *d* + 1 scalar measures, the Minkowski functionals [35]. In this sense, they form a complete set of such global shape descriptors. Although Hadwiger’s theorem is stated for convex bodies, the resulting Minkowski functionals extend, by additivity and continuity, from convex sets to finite unions and to sets with sufficiently regular boundaries, including the non-convex excursion sets of our smoothed fields [43]. In the context of two-dimensional fields, three Minkowski functionals *W*_0_, *W*_1_, and *W*_2_ are defined for a given excursion set *Q* (Figure 1e) as:

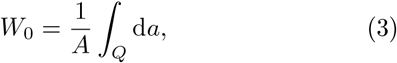

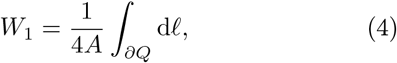

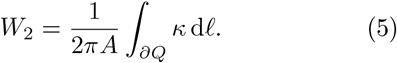

Here, *A* is the total area spanned by the considered gene distribution, and d*a* represents the infinitesimal area element. *∂Q* denotes the boundary of the excursion set and d*ℓ* is the infinitesimal length element along the boundary. *κ* is the local curvature of the boundary. Their discretised estimators, used in practice for numerical evaluation on pixelised excursion sets, can be found in [44].

The first Minkowski functional, *W*_0_, quantifies the fraction of the area occupied by the excursion set, directly capturing how the gene expression distribution extends at a given level set. The second functional, *W*_1_, measures the boundary length of the excursion set. It probes how intricate and fragmented the expression domains are. For the same covered area, a highly speckled pattern with many small patches will have a larger *W*_1_ than a single compact patch. The third Minkowski functional, *W*_2_, is proportional to the Euler characteristic. In two dimensions, this descriptor increases when the excursion set is split into many disconnected regions and decreases when the regions contain holes. It quantifies whether the distribution forms isolated clusters, a percolating network, or perforated “ring-like” structures. It therefore captures the topological organisation of expression domains.

Importantly, Minkowski functionals possess intrinsic mathematical properties that directly address common sources of batch and sample variability. Indeed, because they are invariant under translation, rotation, and reflection [35, 43], the same spatial pattern yields identical Minkowski functional values, whether it has been shifted on the slide, rotated during imaging or flipped. This means that differences in how each tissue section was mounted, masked, or oriented, do not bias comparisons between genes of the same or different samples. Moreover, Minkowski functionals are additive [35, 43]: the measure of the union of disjoint spatial regions equals the sum of the measures of each region. This additivity allows measures to be combined over disjoint regions within a given sample. When replicate samples are available, we separately compute Minkowski characteristics for each sample and then average them to obtain a condition-level representation while reducing sample-specific noise.

To build intuition for the behaviour of the Minkowski functionals, we apply them to a set of controlled, synthetic “toy” spatial patterns in Supplementary Materials section S1.

### 4.3 Computing the Minkowski Tensor

A common limitation of many current ST methods is the limited explicit treatment of distribution anisotropy [4, 45, 12, 46], i.e. directional structure and elongation patterns that are difficult to analyse and model. While Minkowski functionals capture a broad range of spatial properties, they are scalar valuations and are thus insensitive to directionality. Consequently, tensorial extensions of Minkowski functionals have been introduced to consider anisotropy [36, 47, 48, 22]. In our context, these tensorial extensions can be used to explicitly encode directional information of gene expression spatial patterns.

Among these tensors, we focus on the rank-2 Minkowski tensor 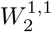. For an excursion set *Q* with boundary *∂Q* and gene distribution area *A*, we define

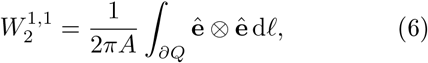

where ê is the unit tangent vector to the boundary *∂Q*, ⊗ denotes the tensor product, and d*ℓ* is the infinitesimal line element along the boundary (see [44] for its discretised version). Like the Minkowski functionals, 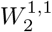 is an additive and translationally invariant Minkowski valuation [22]. Because the tensor is covariant under global rotations, its eigenvalues, and any scalar derived from them, do not depend on the absolute orientation of the pattern.

Consequently, we summarise the information carried by 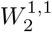 through a scalar anisotropy index. For each connected excursion region, we diagonalise 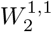 and denote its ordered eigenvalues by *λ*_1_ ≥ *λ*_2_. The dimensionless ratio *λ*_2_*/λ*_1_ ∈ (0, 1] is invariant under translations, global rotations and global scale transformations of the field. It quantifies how isotropic or elongated the region is: values close to 1 indicate nearly isotropic, disk-like shapes, whereas smaller values correspond to increasingly anisotropic, elongated domains [49, 44]. We then define the Minkowski tensor anisotropy index *β* as the mean eigenvalue ratio over all connected regions identified by the friends-of-friends algorithm,

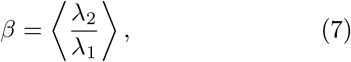

where ⟨·⟩ denotes the average over regions. Throughout this manuscript, we use *β* as our Minkowskitensor–based summary of directional structure.

As for the scalar Minkowski functionals, when multiple replicate samples are available for the same condition, the anisotropy index *β* can be averaged across replicates at each level set.

In summary, a given gene density field generates a sequence of *N*_*ℓ*_ excursion sets (Figure 1d). We get one excursion set per level set. For each excursion set, we compute the four MCs (*W*_0_, *W*_1_, *W*_2_, *β*). These descriptors describe complementary aspects of the spatial geometry. Concatenating the four MCs across all level sets yields a 4 × *N*_*ℓ*_-dimensional representation of the gene spatial _rganization. We refer to this representation as the gene’s Minkowski profile (Figure 1e).

### 4.4 Robust Scaling of Minkowski Profiles

At this stage of the framework, each gene distribution is represented by a Minkowski profile, i.e., a vector composed of the four MCs computed over a series of level sets. Because these four MCs have different physical units and can span different orders of magnitude, we rescale them before downstream analyses.

For each MC *k* ∈ {*W*_0_, *W*_1_, *W*_2_, *β*}, we pool all its values across genes, samples, and level sets. Denoting by *k*_*g,s,ℓ*_ the value of the characteristic *k* for gene *g*, sample *s*, and level set *ℓ*, and by

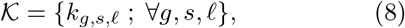

the pooled collection of values for the characteristic *k*, we define the robust-scaled value as

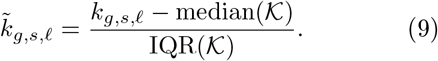

This procedure is applied independently to *W*_0_, *W*_1_, *W*_2_, and *β*. It places all MCs on a comparable scale and thereby prevents any single characteristic from dominating downstream analyses. Importantly, scaling is performed jointly across all the samples considered in a given analysis, as per-sample independent scaling could remove shifts in MC distributions between conditions and attenuate biologically meaningful condition-specific spatial signatures. In contrast to z-score normalization, which relies on the mean and standard deviation and is therefore sensitive to heavy tails and outliers, robust scaling does not assume Gaussianity of MC distributions across genes and samples and is less affected by extreme values.

### 4.5 Covariance Matrix Estimation

Downstream analyses can be performed either with or without accounting for uncertainty. Euclidean distances between Minkowski profiles (Section 4.7) treat each profile as a point estimate in a vector space and do not consider covariance estimation. By contrast, uncertainty-aware analyses require, for each gene in each sample, an estimate of the covariance matrix of its Minkowski profile. This covariance matrix quantifies uncertainties and correlations between the 4 × *N*_*ℓ*_ vector components obtained by concatenating (*W*_0_, *W*_1_, *W*_2_, *β*) across all level sets. It is then used to compute uncertainty-aware 2-Wasserstein distances between Minkowski profiles.

We estimate this covariance matrix using a Monte Carlo resampling procedure. Let 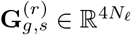 denote the Minkowski profile after robust scaling obtained for gene *g* in sample *s* from Monte Carlo realization *r*, with *r* = 1, …, *N*_real_. The goal of the resampling procedure is to generate alternative stochastic realisations of the same underlying gene spatial organisation, in order to quantify variability induced by finite transcript counts, also known as shot noise, rather than by biological heterogeneity.

To this end, we first infer a continuous density field directly from the original discrete transcript positions using bilinear interpolation. This interpolation yields a grid of continuous density values. To generate synthetic discrete realisations, we then convert this field into integer transcript counts per pixel by Poisson sampling, while preserving the expected global density level. We use Poisson sampling as a baseline of finite-count variability in our resampling procedure. Next, we assign simulated transcript coordinates within each pixel using a bilinear assignment procedure. To this end, we adapt to two dimensions the methodology described in [50], which ensures continuity between adjacent pixels, computational efficiency, and consistency between the interpolation and assignment steps.

This bilinear interpolation step belongs specifically to the Monte Carlo resampling procedure and should not be confused with the Gaussian smoothing used to compute the MCs (Section 4.1). Indeed, these two procedures serve different purposes: the Gaussian smoothing defines the continuous fields on which the MCs are measured, whereas the bilinear interpolation is used here only to construct a simple and tractable resampling scheme. Bilinear interpolation is consistent with the subsequent bilinear assignment of simulated transcript positions, whereas we are not aware of an equally simple semi-analytical construction allowing simulated transcript positions to be reassigned in a way that is exactly consistent with a Gaussian kernel.

Repeating this resampling procedure *N*_real_ times yields an ensemble of synthetic transcript distributions for each gene in each sample. For each realisation *r*, we then compute the full Minkowski profile 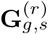 using the same pipeline as for the original data, including Gaussian smoothing, level set construction, and computation of (*W*_0_, *W*_1_, *W*_2_, *β*).

We define the empirical mean profile

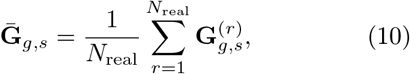

and estimate the covariance matrix of the Minkowski profile by the unbiased empirical estimator

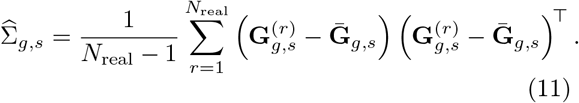

Thus, each gene in each sample is associated with one covariance matrix

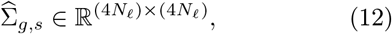

which captures uncertainties and correlations between all Minkowski characteristics and level sets composing its Minkowski profile.

In `minkiPy`, the number of Monte Carlo realisations *N*_real_ is tied to the dimensionality of the Minkowski profile in order to stabilise covariance estimation. For each gene, the profile has dimension *d* = 4*N*_*ℓ*_. By default, `minkiPy` sets

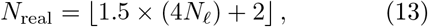

so that the number of Monte Carlo samples exceeds the minimum *d* + 1 required for a full-rank empirical covariance matrix. This choice reduces the risk of rank-deficient or ill-conditioned covariance estimates while keeping the computational cost under control.

### 4.6 Averaging Samples

Datasets may include several samples from the same biological condition, for instance biological replicates, adjacent slices, or different regions of interest of a given sample. In `minkiPy`, the gene’s Minkowski profiles can be averaged. In the FSHD MERFISH case we constructed condition-level averaged samples from biological replicates belonging to the same conditions.

Let G_*g,s*_ denote the robust-scaled Minkowski profile of gene *g* in sample *s*, with dimension *d* = 4*N*_*ℓ*_, and let 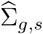 denote its Monte Carlo covariance matrix, if provided. For a given condition *c* represented by *K* samples *s* ∈ *c*, we define the condition-level profile as the simple arithmetic mean

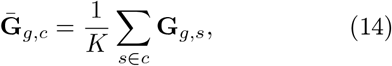

treating the individual samples as independent realisations of the same underlying condition-level spatial pattern. Under this independence assumption, the covariance of the mean profile is

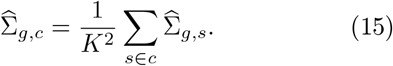

The resulting averaged profiles and covariances define a condition-level synthetic sample for each group of replicates. This averaged sample is then used in downstream analyses to represent the biological signal associated with the condition. Sample-specific noise is reduced by the averaging process.

### 4.7 Metric Selection and Gene Distance Computation

To compare the spatial pattern of genes (different genes from the same or different samples or same gene from different samples), we need a metric that quantifies the distances between their Minkowski profiles. Each profile is represented as a point in a multidimensional feature space, which is shared by all genes. The dimension of this space is 4*N*_*ℓ*_, which corresponds to the four MCs evaluated across *N*_*ℓ*_ level sets. Optionally, each profile can additionally be associated with a covariance matrix estimated from Monte Carlo realisations. Our goal is to compute distances between profiles in a way that respects both their location in the feature space, as well as their uncertainty if provided.

When no covariance matrix is provided, we quantify the distances between Minkowski profiles using the standard Euclidean distance. If 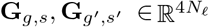 denote the robust-scaled Minkowski profiles of genes *g* in sample *s* and *g*′ in sample *s*′, their distance is

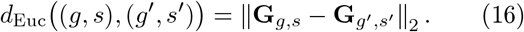

This standard Euclidean distance metric treats profiles as point estimates and ignores uncertainty, but provides a simple and computationally efficient baseline for comparing gene spatial patterns.

When covariance information is available and uncertainty-aware comparisons are desired, we use the Gaussian closed form of the 2-Wasserstein distance [51]. This Gaussian closed form is justified by the empirical observation that Monte Carlo resampling yields distributions of Minkowski profiles that are well approximated by multivariate Gaussians (Figure S21). If a gene profile is represented by 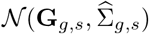 and another by 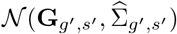, their squared 2-Wasserstein distance is

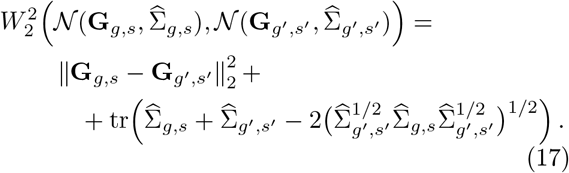

This expression is the closed-form solution of the optimal transport problem between Gaussian measures in the 2-Wasserstein geometry. The squared 2-Wasserstein distance reduces to the squared Euclidean distance (Eq. 16) between profile means when 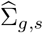 and 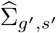 vanish, and can therefore be viewed as a covariance-aware extension of the Euclidean metric.

To reduce computation time on large datasets, we also provide an option to approximate each full covariance matrix by its diagonal. This replaces the full Gaussian 2-Wasserstein distance by a diagonal-covariance approximation that neglects correlations between MCs and between level sets, thereby reducing the cost of distance computation. We found that using only diagonal covariances yields 2-Wasserstein distances that are nearly identical to those obtained with the full covariances (Figure S22).

Using either the Euclidean distance or the covariance-aware 2-Wasserstein distance (with full or diagonal-approximated covariances), we compute distances between all available gene Minkowski profiles. These computations may involve different genes within the same sample, the same gene across different samples, or different genes across different samples. The result is a symmetric distance matrix

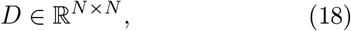

where *N* denotes the total number of Minkowski profiles under consideration. This matrix constitutes the basic output of the distance computation step and serves as input for downstream analyses.

### 4.8 Similarity Graph Construction and Community Detection

To identify communities of genes sharing similar spatial organisation, we construct a similarity graph in which genes are represented as nodes and edges encode the similarity between the nodes based on their distances in the Minkowski profile space.

To construct the similarity graph, we transform the distance matrix *D* ∈ ℝ^*N×N*^ into a dense similarity matrix *S* ∈ ℝ^*N×N*^ using a Gaussian radial basis function (RBF) kernel,

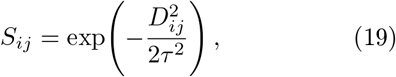

where *τ* controls the length scale of the RBF kernel. By default, we set *τ* to the median of non-zero distances, which yields a data-driven bandwidth and limits sensitivity to outliers.

Because the RBF kernel is a monotonic transform of the distance matrix, the ordering of pairwise relationships is preserved. We then sparsify *S* by constructing a weighted *k*-nearest-neighbour graph, in which edges connect nodes with highest affinities. To guarantee global connectivity and avoid isolated components, we additionally augment this graph with edges from the minimum spanning tree computed on *D*.

Community detection is then performed on the resulting sparse weighted *k*-nearest-neighbour graph using the Leiden algorithm [52]. Parameters selected for the use-cases are provided in the parameter choices Section 4.11 and in the figure captions.

### 4.9 Gene-level and Sample-level Low Dimensional Embeddings

We construct low-dimensional embeddings at two complementary levels: the gene level and the sample level. In both cases, embeddings are built either using directly the Minkowski profile space or using distance matrices derived from the Minkowski profiles.

At the gene level, each observation corresponds to the Minkowski profile of one gene in one sample (or one gene in one condition-averaged sample, when such averaging is used). For gene-level embedding based on Minkowski profiles directly, we stack all robust-scaled Minkowski profiles 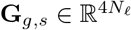 into a data matrix

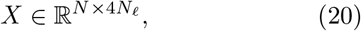

where *N* is the total number of profiles under consideration. We apply principal component analysis (PCA) to *X* to obtain a linear embedding that captures dominant modes of variation in Minkowski profile space. Uniform Manifold Approximation and Projection (UMAP) is also applied to *X*, using the Euclidean metric in Minkowski feature space, to obtain a non-linear embedding that preserves local neighbourhood structure in this feature space.

For gene-level embedding based on distances computed between gene Minkowski profiles, we use the distance matrix

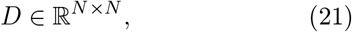

defined in Section 4.7. Depending on the analysis, *D* may contain either Euclidean distances or covariance-aware 2-Wasserstein distances between Minkowski profiles. We apply classical multidimensional scaling (MDS) to *D* using the standard Torgerson–Gower procedure [53]. This yields a two-dimensional Euclidean embedding that best preserves the original distance geometry in a least-squares sense.

The Leiden communities detected from the similarity graph (Section 4.8) are overlaid on the gene-level PCA, UMAP, and MDS embeddings.

For sample-level embeddings, we consider a collection of *F* samples. For sample-level embedding based on Minkowski profiles directly, each sample *s* is represented by a high-dimensional vector obtained by concatenating the robust-scaled Minkowski profiles of all genes of that sample. If *N*_*g*_ denotes the number of genes shared across the samples under consideration, this yields a matrix

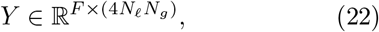

with one row per sample. We perform PCA on *Y* to obtain a linear embedding that summarises global variation in spatial gene organisation across samples. We do not use UMAP at the sample level, because the number of samples is typically too small.

For sample-level embedding based on distances, we define a sample–sample distance matrix

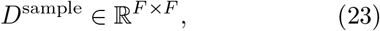

which aggregates gene-level distances between pairs of samples. For each pair of samples (*s, s*′) and each shared gene *g*, let *d*_*s,s*_′ (*g*) denote the distance (Euclidean or 2-Wasserstein) between the Minkowski profiles of the same gene *g* in samples *s* and *s*′. We then compute

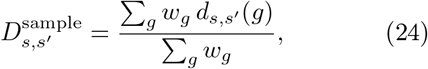

where the sum runs over genes shared by samples *s* and *s*′, and the weights *w*_*g*_ are proportional to the total number of transcripts of gene *g* in the two samples. This transcript-count weighted averaging emphasises genes that are well sampled in both samples and down-weights genes with sparse or noisy measurements. We then apply classical MDS to *D*^sample^ to obtain a two-dimensional embedding.

Throughout this work, we restrict comparisons to genes that are shared across all samples under consideration. This ensures that gene-level and sample-level distance matrices, as well as feature-based embeddings, are defined on a common gene set and remain directly comparable across samples.

### 4.10 Sample-level Differential Gene Expression

We compute normalised gene densities as a simple global measure of gene expression across samples. For each sample *s*, let *N*_*g,s*_ denote the total number of transcripts of gene *g*, and let *A*_*s*_ denote the tissue area covered by the sample. We first compute a raw density per unit area:

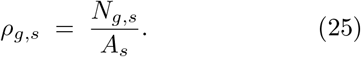

Because total transcript counts,

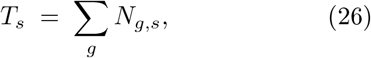

can vary substantially between samples, for example owing to differences in capture efficiency, sequencing depth, or tissue coverage, we apply a simple global rescaling across samples. Let 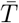 be the mean of *T*_*s*_ over all samples under consideration, and define a sample-specific scale factor 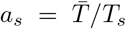. The normalised density is then:

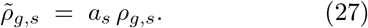

This procedure harmonises the overall density level between samples while preserving within-sample relative differences between genes.

For condition-level averaged samples (Section 4.6), we define the mean density as the arithmetic mean of the normalised densities across all samples belonging to the same condition.

To quantify differential gene expression between two samples *s*_*A*_ and *s*_*B*_, we compute a log_2_ fold change based on the normalised densities. For each gene *g*, we define

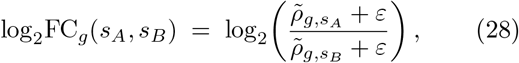

where *ε* is a small pseudo-count added to avoid numerical instability when densities are close to zero. In addition, we report the relative density variation,

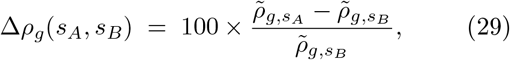

whenever the denominator is non-zero.

These quantities provide gene-level measures of global gene expression change that can be directly compared with spatial metrics derived from Minkowski profiles, such as Euclidean or 2−Wasserstein distances.

### 4.11 Parameter Choices and Filtering Criteria for the FSHD MER-FISH and CRC Visium HD Datasets

For the FSHD MERFISH dataset, Minkowski characteristics are evaluated on 30 linearly spaced level sets (Section 2.2); for the CRC Visium HD dataset, Minkowski characteristics are evaluated on 20 linearly spaced level sets (Section 2.3). A spatial grid resolution of 20 *µ*m is used for all samples in both datasets. Comparable results are obtained with resolutions of 5, 10, 20, or 50 *µ*m.

For the FSHD MERFISH dataset, Minkowski profiles are computed on the complete panel of probed genes. For the CRC Visium HD dataset, Minkowski profiles are computed only for genes with at least 100 detected transcripts within a given sample. In addition, when focusing on a given set of samples, we retain only (i) the intersection of genes present in all samples and (ii) genes with at least 2000 detected transcripts in at least one sample.

For both the FSHD MERFISH and the CRC Visium HD datasets, Monte Carlo covariance matrices are estimated using

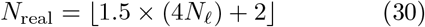

Monte Carlo realisations.

For graph construction and community detection, we use in both datasets a *k*-nearest-neighbour graph augmented with a minimum spanning tree. For the FSHD MERFISH dataset, the graph is built with *k* = 25, and communities are identified with the Leiden algorithm using a resolution parameter of 0.05. For the CRC Visium HD dataset, we use a denser graph with *k* = 200, and Leiden clustering is performed with a resolution parameter of 0.04.

### 4.12 The minkiPy Package

`minkiPy` is an open-source Python package. The source code, documentation and example workflows for `minkiPy` are available at https://github.com/BAUDOTlab/minkiPy. For imaging-based assays, users provide per-transcript (*x, y*) coordinates. For sequencing-based assays, users represent each transcript as a separate row and assign to it the (*x, y*) coordinates of the centroid of the spot in which it was detected. The method otherwise treats all spatial transcriptomics technologies identically.

Users must specify two parameters: the grid resolution and the number of level sets. All other parameters are optional. Defaults are provided for the number of Monte Carlo realisations, the use of diagonal or full covariance matrices in the 2-Wasserstein distance, Leiden clustering settings, and plotting parameters. When the number of Monte Carlo realisations is set to zero, covariance matrices are not estimated and downstream analyses automatically use Euclidean distances between Minkowski profiles.

The package is parallelised with MPI and is designed for CPU architectures. It uses shared-memory data structures to limit RAM usage across workers. Workflows can be launched from the command line with `mpiexec` or from Python scripts and notebooks. The package is available as open-source code, together with documentation, and example workflows.

### 4.13 Computational performance

For the MERFISH FSHD sample `batch4_Control`, containing 37,120,299 transcripts (∼ 2.89 GB input table), the computation took 1086.76 s (∼ 18.1 min), with an estimated peak RAM usage of 5.77 GiB. 30 CPU threads were used.

For the Visium HD CRC sample `P2_CRC`, containing 129,467,679 transcripts (∼ 20.86 GB input table), the computation took 40888.30 s (∼ 11.4 h), with an estimated peak RAM usage of 40.57 GiB. 30 CPUs were used.

## Data and Code Availability

`minkiPy` is available at https://github.com/BAUDOTlab/minkiPy. The datasets analysed in this study are publicly available from the original publications cited in the manuscript. Documentation, source code, and reproducibility notebooks enabling the reproduction of the main analyses and results are provided through the associated GitHub repository.

## Acknowledgements

The authors thank Lionel Spinelli, Franck Picard, Marielle Péré, Bastien Chassagnol and Morgane Térézol for carefully reading the manuscript and providing useful feedback. The authors also thank Julien Bel and Clément Moroldo for useful discussions.

## References

[1] Patrik L Ståhl, Fredrik Salmén, Sanja Vickovic, Anna Lundmark, José Fernández Navarro, Jens Magnusson, Stefania Giacomello, Michaela Asp, Jakub O Westholm, Mikael Huss, et al. Visualization and analysis of gene expression in tissue sections by spatial transcriptomics. Science, 353(6294):78–82, 2016.

[2] Kok Hao Chen, Alistair N Boettiger, Jeffrey R Moffitt, Siyuan Wang, and Xiaowei Zhuang. Spatially resolved, highly multiplexed rna profiling in single cells. Science, 348(6233):aaa6090, 2015.

[3] Kristen R Maynard, Leonardo Collado-Torres, Lukas M Weber, Cedric Uytingco, Brianna K Barry, Stephen R Williams, Joseph L Catallini, Matthew N Tran, Zachary Besich, Madhavi Tippani, et al. Transcriptome-scale spatial gene expression in the human dorsolateral prefrontal cortex. Nature neuroscience, 24(3):425–436, 2021.

[4] Lambda Moses and Lior Pachter. Museum of spatial transcriptomics. Nature methods, 19(5):534–546, 2022.

[5] Zhiyuan Yuan, Fangyuan Zhao, Senlin Lin, Yu Zhao, Jianhua Yao, Yan Cui, Xiao-Yong Zhang, and Yi Zhao. Benchmarking spatial clustering methods with spatially resolved transcriptomics data. Nature Methods, 21(4):712–722, 2024.

[6] Erick Armingol, Hratch M Baghdassarian, and Nathan E Lewis. The diversification of methods for studying cell–cell interactions and communication. Nature Reviews Genetics, 25(6):381–400, 2024.

[7] Zhijian Li, Zain M. Patel, Dongyuan Song, Sai Nirmayi Yasa, Robrecht Cannoodt, Guanao Yan, Jingyi Jessica Li, and Luca Pinello. Systematic benchmarking of computational methods to identify spatially variable genes. Genome Biology, 26(1):285, 2025.

[8] Brendan F Miller, Dhananjay Bambah-Mukku, Catherine Dulac, Xiaowei Zhuang, and Jean Fan. Characterizing spatial gene expression heterogeneity in spatially resolved single-cell transcriptomic data with nonuniform cellular densities. Genome research, 31(10):1843–1855, 2021.

[9] David DeTomaso and Nir Yosef. Hotspot identifies informative gene modules across modalities of single-cell genomics. Cell systems, 12(5):446–456, 2021.

[10] Duy Pham, Xiao Tan, Brad Balderson, Jun Xu, Laura F Grice, Sohye Yoon, Emily F Willis, Minh Tran, Pui Yeng Lam, Arti Raghubar, et al. Robust mapping of spatiotemporal trajectories and cell–cell interactions in healthy and diseased tissues. Nature communications, 14(1):7739, 2023.

[11] Ron Zeira, Max Land, Alexander Strzalkowski, and Benjamin J Raphael. Alignment and integration of spatial transcriptomics data. Nature Methods, 19(5):567–575, 2022.

[12] Kalen Clifton, Manjari Anant, Gohta Aihara, Lyla Atta, Osagie K Aimiuwu, Justus M Kebschull, Michael I Miller, Daniel Tward, and Jean Fan. Stalign: Alignment of spatial transcriptomics data using diffeomorphic metric mapping. Nature communications, 14(1):8123, 2023.

[13] Patrick CN Martin, Wenqi Wang, Hyobin Kim, Henrietta Holze, Paul B Fisher, Arturo P Saavedra, Robert A Winn, Esha Madan, Rajan Gogna, and Kyoung Jae Won. Multi-scale and multi-context interpretable mapping of cell states across heterogeneous spatial samples. Nature Communications, 16(1):7814, 2025.

[14] Wei Liu, Xu Liao, Ziye Luo, Yi Yang, Mai Chan Lau, Yuling Jiao, Xingjie Shi, Weiwei Zhai, Hongkai Ji, Joe Yeong, et al. Probabilistic embedding, clustering, and alignment for integrating spatial transcriptomics data with precast. Nature communications, 14(1):296, 2023.

[15] Wei Liu, Xu Liao, Yi Yang, Huazhen Lin, Joe Yeong, Xiang Zhou, Xingjie Shi, and Jin Liu. Joint dimension reduction and clustering analysis of single-cell rna-seq and spatial transcriptomics data. Nucleic acids research, 50(12):e72–e72, 2022.

[16] Peiying Cai, Mark D Robinson, and Simone Tiberi. Despace: spatially variable gene detection via differential expression testing of spatial clusters. Bioinformatics, 40(2):btae027, 2024.

[17] Fei Qin, Xizhi Luo, Qing Lu, Bo Cai, Feifei Xiao, and Guoshuai Cai. Spatial pattern and differential expression analysis with spatial transcriptomic data. Nucleic Acids Research, 52(21):e101–e101, 2024.

[18] Hermann Minkowski. Volumen und oberfläche. Mathematische Annalen, 57:447–495, 1903.

[19] K. R. Mecke, T. Buchert, and H. Wagner. Robust morphological measures for large scale structure in the universe. Astron. Astrophys., 288:697–704, 1994.

[20] Jens Schmalzing and Krzysztof M Górski. Minkowski functionals used in the morphological analysis of cosmic microwave background anisotropy maps. Monthly Notices of the Royal Astronomical Society, 297(2):355–365, 1998.

[21] Jens Schmalzing, Stefan Gottlöber, Anatoly A Klypin, and Andrey V Kravtsov. Quantifying the evolution of higher order clustering. Monthly Notices of the Royal Astronomical Society, 309(4):1007–1016, 1999.

[22] Gerd E Schröder-Turk, Walter Mickel, Sebastian C Kapfer, Fabian M Schaller, Boris Breidenbach, Daniel Hug, and Klaus Mecke. Minkowski tensors of anisotropic spatial structure. New Journal of Physics, 15(8):083028, 2013.

[23] Timothy J Larkin, Holly C Canuto, Mikko I Kettunen, Thomas C Booth, De-En Hu, Anant S Krishnan, Sarah E Bohndiek, André A Neves, Charles McLachlan, Michael P Hobson, et al. Analysis of image heterogeneity using 2d minkowski functionals detects tumor responses to treatment. Magnetic resonance in medicine, 71(1):402–410, 2014.

[24] Marconi Barbosa, Riccardo Natoli, Kriztina Valter, Jan Provis, and Ted Maddess. Integral-geometry characterization of photobiomodulation effects on retinal vessel morphology. Biomedical Optics Express, 5(7):2317–2332, 2014.

[25] Anne Savage, Elad Katz, Alistair Eberst, Ruth E Falconer, Alasdair Houston, David J Harrison, and James Bown. Characterising the tumour morphological response to therapeutic intervention: an ex vivo model. Disease models & mechanisms, 6(1):252–260, 2013.

[26] Lujia Chen, Xiangduo Kong, Kevin G Johnston, Ali Mortazavi, Todd C Holmes, Zhiqun Tan, Kyoko Yokomori, and Xiangmin Xu. Single-cell spatial transcriptomics reveals a dystrophic trajectory following a developmental bifurcation of myoblast cell fates in facioscapulohumeral muscular dystrophy. Genome Research, 34(5):665–679, 2024.

[27] Michelli Faria de Oliveira, Juan Pablo Romero, Meii Chung, Stephen R Williams, Andrew D Gottscho, Anushka Gupta, Susan E Pilipauskas, Seayar Mohabbat, Nandhini Raman, David J Sukovich, et al. High-definition spatial transcriptomic profiling of immune cell populations in colorectal cancer. Nature Genetics, pages 1–12, 2025.

[28] Liangbo Zhao, Xiao Sun, Chenying Hou, Yanmei Yang, Peiwen Wang, Zhaoyuan Xu, Zhenzhen Chen, Xiangrui Zhang, Guanghua Wu, Hong Chen, et al. Cpne7 promotes colorectal tumorigenesis by interacting with nono to initiate zfp42 transcription. Cell Death & Disease, 15(12):896, 2024.

[29] Guido Giordano, Pietro Parcesepe, Mario Rosario D’Andrea, Luigi Coppola, Tania Di Raimo, Andrea Remo, Erminia Manfrin, Claudia Fiorini, Aldo Scarpa, Carla Azzurra Amoreo, et al. Jak/stat5-mediated subtype-specific lymphocyte antigen 6 complex, locus g6d (ly6g6d) expression drives mismatch repair proficient colorectal cancer. Journal of Experimental & Clinical Cancer Research, 38(1):28, 2019.

[30] Graeme I Murray, Margaret E Duncan, Pauline O’Neil, William T Melvin, and John E Fothergill. Matrix metalloproteinase–1 is associated with poor prognosis in colorectal cancer. Nature medicine, 2(4):461–462, 1996.

[31] Jing Yu, Zhen He, Xiaowen He, Zhanhao Luo, Lei Lian, Baixing Wu, Ping Lan, and Haitao Chen. Comprehensive analysis of the expression and prognosis for mmps in human colorectal cancer. Frontiers in Oncology, 11:771099, 2021.

[32] Chunjie Xu, Guangang Tian, Chunhui Jiang, Hanbing Xue, Manzila Kuerbanjiang, Longci Sun, Lei Gu, Hong Zhou, Ye Liu, Zhigang Zhang, et al. Nptx2 promotes colorectal cancer growth and liver metastasis by the activation of the canonical wnt/β-catenin pathway via fzd6. Cell death & disease, 10(3):217, 2019.

[33] John W Schoggins, Sam J Wilson, Maryline Panis, Mary Y Murphy, Christopher T Jones, Paul Bieniasz, and Charles M Rice. A diverse range of gene products are effectors of the type i interferon antiviral response. Nature, 472(7344):481–485, 2011.

[34] Hume A. Feldman, Nick Kaiser, and John A. Peacock. Power spectrum analysis of threedimensional redshift surveys. Astrophys. J., 426:23–37, 1994.

[35] Hugo Hadwiger. Vorlesungen über inhalt, oberfläche und isoperimetrie, volume 93. Springer-Verlag, 2013.

[36] Semyon Alesker. Continuous rotation invariant valuations on convex sets. Annals of Mathematics, 149(3):977–1005, 1999.

[37] Tongxuan Lv, Ying Zhang, Mei Li, Qiang Kang, Shuangsang Fang, Yong Zhang, Susanne Brix, and Xun Xu. Eags: efficient and adaptive gaussian smoothing applied to high-resolved spatial transcriptomics. GigaScience, 13:giad097, 2024.

[38] Aoxiang Jiang, Wei Liu, Wenjuan Fang, Baojiu Li, Cristian Barrera-Hinojosa, and Yufei Zhang. Minkowski functionals of large-scale structure as a probe of modified gravity. Physical Review D, 109(8):083537, 2024.

[39] Hubert Mantz, Karin Jacobs, and Klaus Mecke. Utilizing minkowski functionals for image analysis: a marching square algorithm. Journal of Statistical Mechanics: Theory and Experiment, 2008(12):P12015, 2008.

[40] Christine S Botzler, J Snigula, R Bender, and U Hopp. Finding structures in photometric redshift galaxy surveys: an extended friends-of-friends algorithm. Monthly Notices of the Royal Astronomical Society, 349(2):425–439, 2004.

[41] Jens Schmalzing and Thomas Buchert. Beyond genus statistics: a unifying approach to the morphology of cosmic structure. The Astrophysical Journal, 482(1):L1, 1997.

[42] Martin Kerscher. Statistical analysis of large-scale structure in the universe. In Statistical Physics and Spatial Statistics: The art of analyzing and modeling spatial structures and pattern formation, pages 36–71. Springer, 2000.

[43] Klaus R Mecke. Additivity, convexity, and beyond: applications of minkowski functionals in statistical physics. In Statistical Physics and Spatial Statistics: The art of analyzing and modeling spatial structures and pattern formation, pages 111–184. Springer, 2000.

[44] Stephen Appleby, Pravabati Chingangbam, Changbom Park, Sungwook E Hong, Juhan Kim, and Vidhya Ganesan. Minkowski tensors in two dimensions: Probing the morphology and isotropy of the matter and galaxy density fields. The Astrophysical Journal, 858(2):87, 2018.

[45] Zhenqin Wu, Ayano Kondo, Monee McGrady, Ethan AG Baker, Benjamin Chidester, Eric Wu, Maha K Rahim, Nathan A Bracey, Vivek Charu, Raymond J Cho, et al. Discovery and generalization of tissue structures from spatial omics data. Cell Reports Methods, 4(8), 2024.

[46] Jose Camacho, Michael Sorochan Armstrong, Luz Garcia-Martinez, Caridad Diaz, and Carolina Gomez-Llorente. Single-cell spatial (scs) omics recent developments in data analysis. TrAC Trends in Analytical Chemistry, page 118109, 2024.

[47] Claus Beisbart, Robert Dahlke, Klaus Mecke, and Herbert Wagner. Vector-and tensor-valued descriptors for spatial patterns. In Morphology of Condensed Matter: Physics and Geometry of Spatially Complex Systems, pages 238–260. Springer, 2002.

[48] Gerd E Schröder-Turk, Sebastian Kapfer, Boris Breidenbach, Claus Beisbart, and Klaus Mecke. Tensorial minkowski functionals and anisotropy measures for planar patterns. Journal of microscopy, 238(1):57–74, 2010.

[49] Vidhya Ganesan and Pravabati Chingangbam. Tensor minkowski functionals: first application to the cmb. Journal of Cosmology and Astroparticle Physics, 2017(06):023, 2017.

[50] Philippe Baratta, Julien Bel, Sylvain Gouyou Beauchamps, and Carmelita Carbone. Covmos: a new monte carlo approach for galaxy clustering analysis. Astronomy & Astrophysics, 673:A1, 2023.

[51] Matthias Gelbrich. On a formula for the l2 wasserstein metric between measures on euclidean and hilbert spaces. Mathematische Nachrichten, 147(1):185–203, 1990.

[52] Vincent A Traag, Ludo Waltman, and Nees Jan Van Eck. From louvain to leiden: guaranteeing well-connected communities. Scientific reports, 9(1):1–12, 2019.

[53] Ingwer Borg and Patrick Groenen. Classical Scaling, pages 207–212. Springer New York, New York, NY, 1997.

