## Supplementary Materials for "Differential Analysis of Gene Spatial Organisation with Minkowski Functionals and Tensors"

*Philippe Baratta*[1], *Paul Villoutreix*[1], *Anaïs Baudot*[1,2]
[1]Aix Marseille Univ, INSERM, MMG, Turing Centre for Living Systems, Marseille, France
[2]CNRS, Marseille, France

#### S1 Illustration of Minkowski Characteristics Using Synthetic Toy Models

To provide intuition for the behaviour of the Minkowski characteristics, we generated eight synthetic point-pattern distributions within a common elliptical tissue mask. These toy models span four broad categories: isotropic kernels (Gaussian and cosine), anisotropic kernels (directionally stretched cosine and Gaussian), concentric shell patterns, and a power-law Gaussian random field. Each distribution was sampled at high density (100,000 points) and processed using the `minkiPy` pipeline: the continuous transcript density field was thresholded at multiple level sets, and the three scalar Minkowski functionals $W_0$, $W_1$ and $W_2$ were computed together with the fourth characteristic, the anisotropy index $\beta$ (derived from the Minkowski tensor) (Supplementary Figure S20). For all toy-model analyses, we used a grid resolution of 20 $\mu$m, evaluated the Minkowski characteristics on 50 linearly spaced level sets, and estimated covariances using 302 Monte Carlo realisations.

#### S2 Relationship between classical MDS and PCA

Classical multidimensional scaling (MDS) applied to the 2-Wasserstein distance matrix yields embeddings that are visually indistinguishable from those obtained by principal component analysis (PCA) applied directly to the Minkowski profiles. Classical MDS reconstructs Euclidean coordinates from a distance matrix derived from pairwise distances between vectors. When the input distances for MDS are Euclidean, MDS is equivalent to performing PCA on the centred feature matrix [1]. The fact that, in our use cases, MDS and PCA yield visually indistinguishable embeddings suggests that the 2-Wasserstein distances computed are close to Euclidean distances (Supplementary Figure S22b and c). This suggests that the contribution of covariance terms is small relative to the difference between profiles, which is what we observe for the MERFISH FSHD dataset (Supplementary Figure S22b).

Although the Euclidean approximation is very good in practice, covariance estimation remains useful when one aims at a more rigorous description of the geometry. In the Visium HD CRC dataset, Supplementary Figure S22c shows that more than half of the Euclidean distances differ by more than 1% from the diagonal 2-Wasserstein approximation, despite prior filtering of very lowly expressed genes (see Methods, Section 4.11).

#### S3 Supplementary Figures

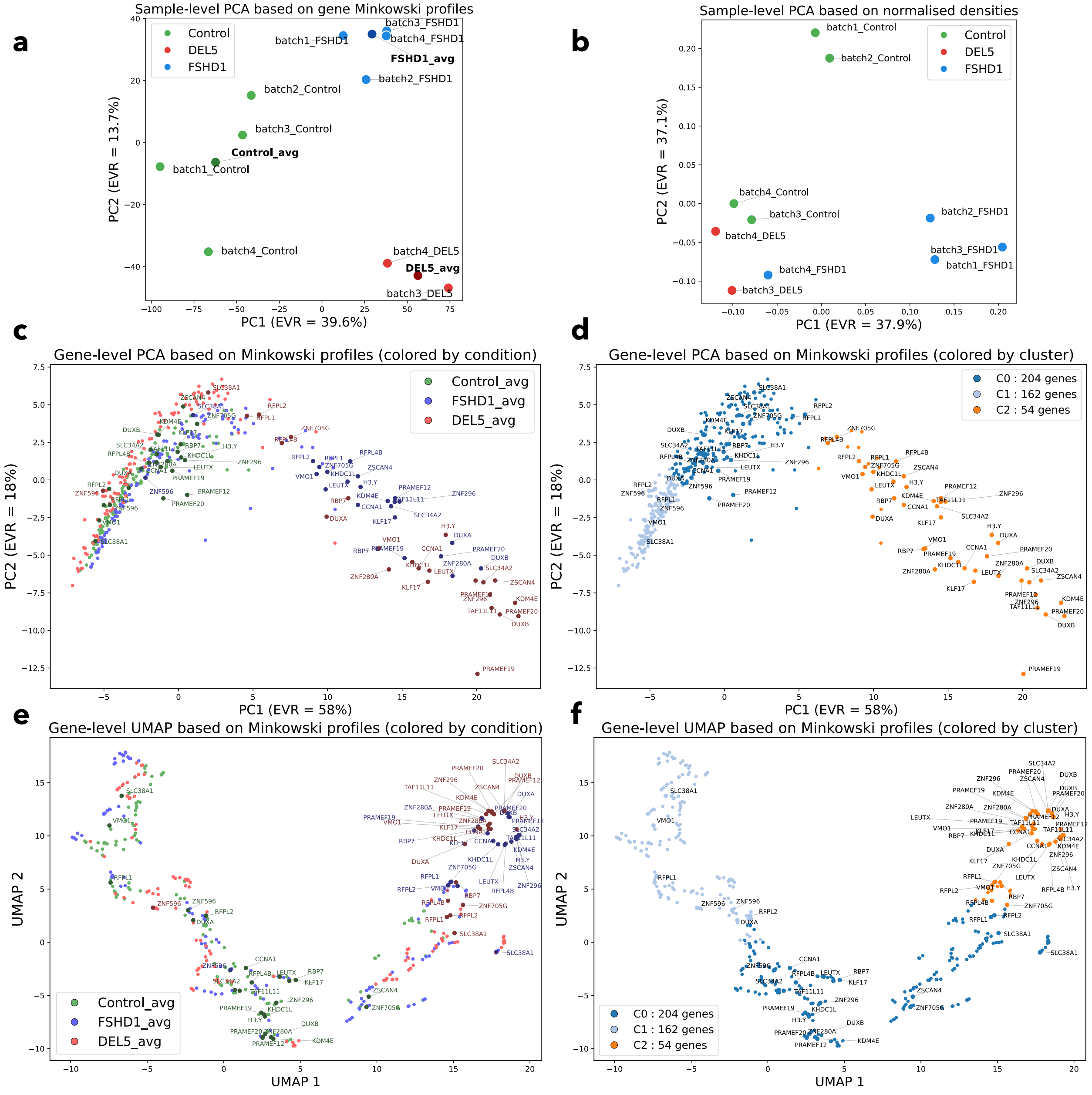


Figure S1: **(a)** PCA of sample-level Minkowski profiles in the FSHD MERFISH dataset. Each point represents one sample, coloured by condition, with darker points indicating the averaged samples for each condition. Axis labels indicate the variance explained by PC1 and PC2. **(b)** PCA of sample-level normalised gene densities. Each point represents one sample, projected from the vector of normalised gene densities and coloured by condition. **(c)** PCA of condition-averaged gene-level Minkowski profiles. Each point represents one gene in one averaged condition; each gene therefore appears three times, once per averaged condition. Points are coloured by condition, and DUX4-target genes are shown with darker markers and labelled. **(d)** The same PCA embedding, coloured by communities identified with the Leiden algorithm. **(e)** UMAP embedding of the same condition-averaged gene-level Minkowski profiles. Each point represents one gene in one averaged condition; each gene therefore appears three times. Points are coloured by condition, and DUX4-target genes are shown with darker markers and labelled. **(f)** The same UMAP embedding, coloured by communities identified with the Leiden algorithm.

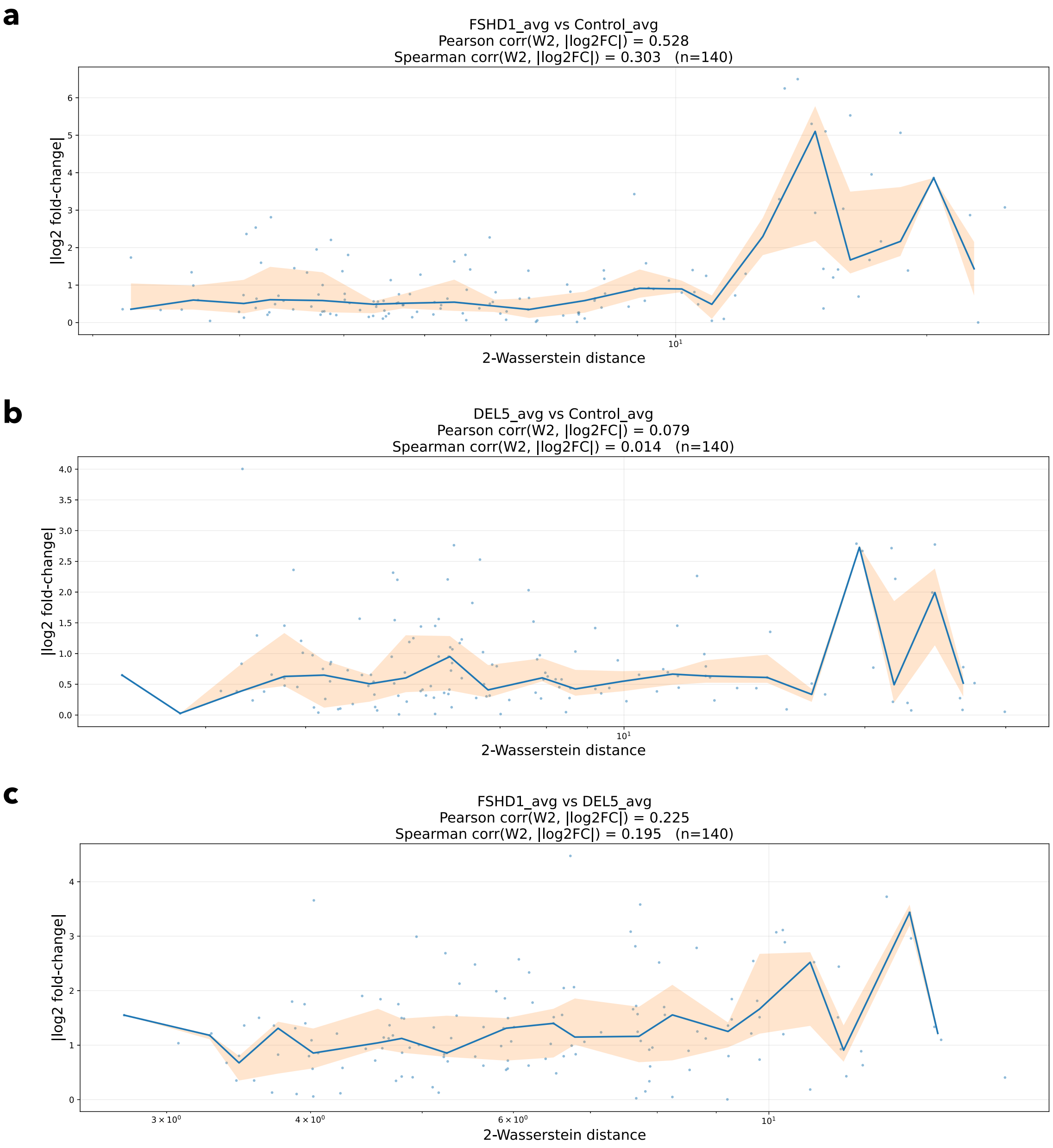


Figure S2: Relationship between spatial reorganisation and differential gene expression in the FSHD MERFISH dataset. Each panel shows, for a pair of averaged conditions, the gene-level 2-Wasserstein distance between Minkowski profiles plotted against the absolute $\log_2$ fold-change computed from normalised gene densities. Each point corresponds to one gene. The solid curve represents the median $|\log_2 \mathrm{FC}|$ within logarithmic bins of 2-Wasserstein distances, and the shaded region indicates the interquartile range. The Pearson and Spearman correlations between the 2-Wasserstein distances and $|\log_2 \mathrm{FC}|$ are reported in each panel. (a) FSHD1_avg vs Control_avg. (b) DEL5_avg vs Control_avg. (c) FSHD1_avg vs DEL5_avg.

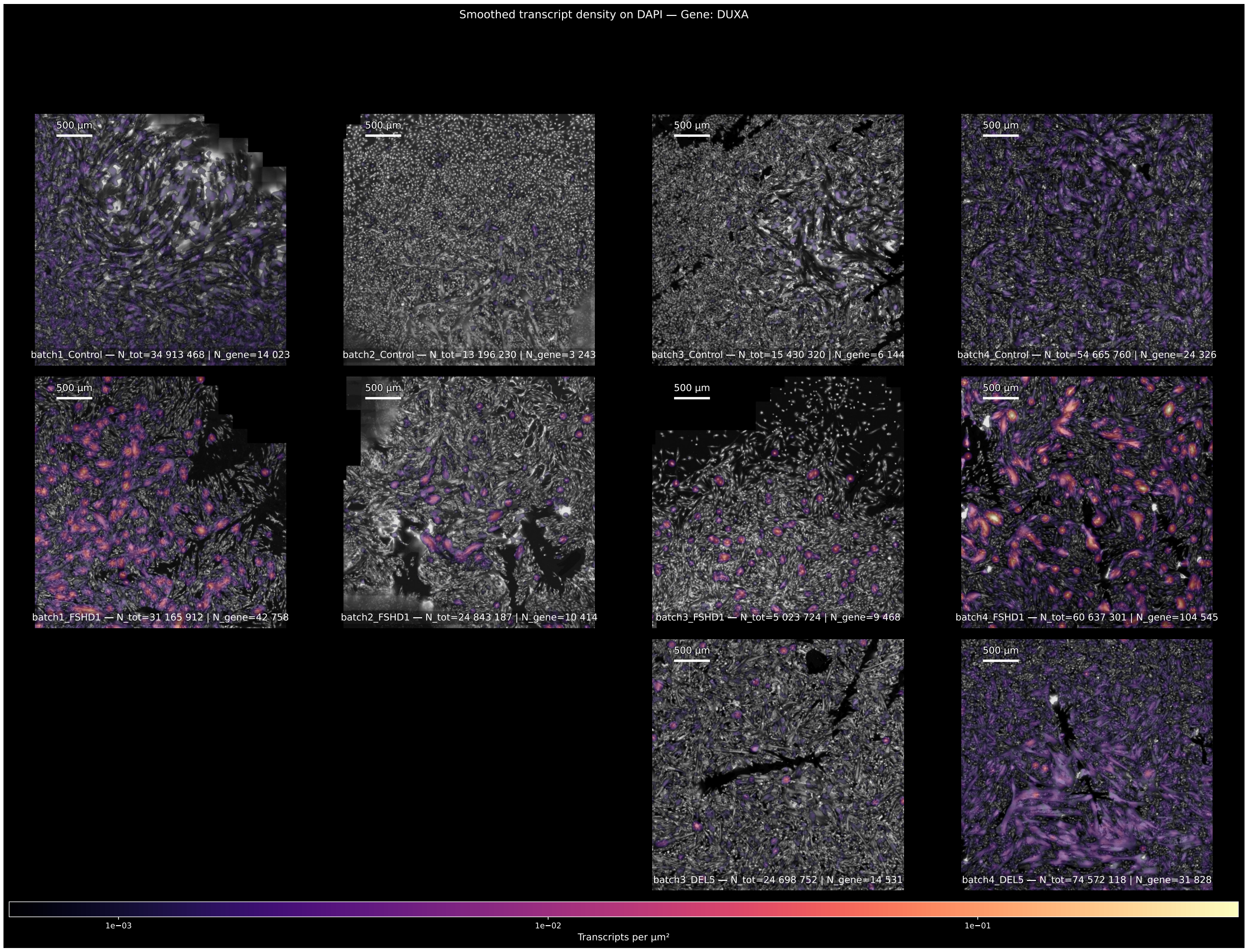


Figure S3: DAPI-normalised spatial density maps for *DUXA* in the FSHD MERFISH dataset. For each sample, the raw *DUXA* transcript point pattern was transformed into a continuous density field using Gaussian smoothing. The resulting smoothed density map is overlaid on the corresponding DAPI image for spatial context. Samples are arranged by condition (Control, FSHD1, and DEL5; rows) and by replicate (columns), with all panels shown at the same spatial scale and using the same crop window and colour scale. A 500 $\mu$m scalebar is shown in each panel.

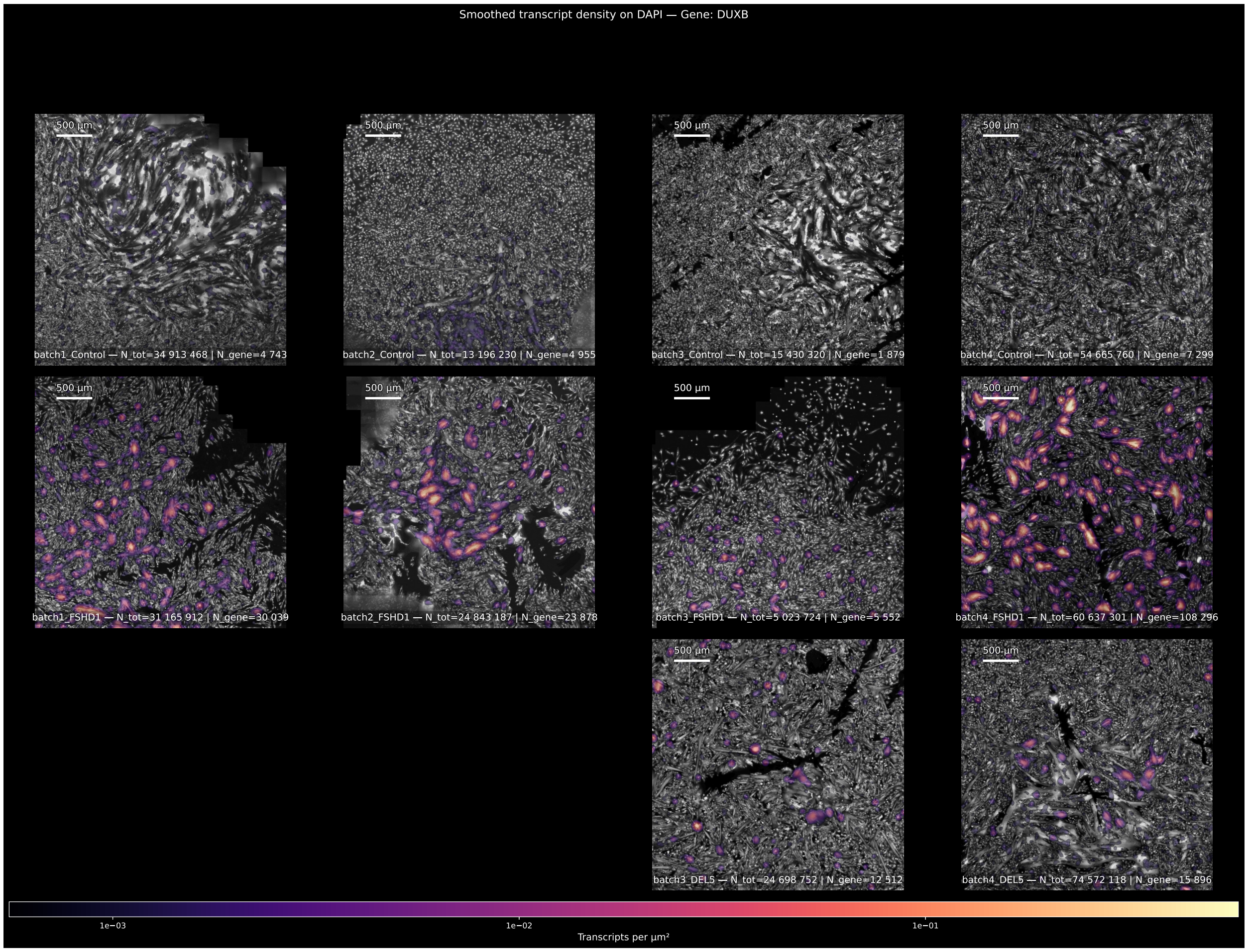


Figure S4: DAPI-normalised spatial density maps for *DUXB* in the FSHD MERFISH dataset. For each sample, the raw *DUXB* transcript point pattern was transformed into a continuous density field using Gaussian smoothing. The resulting smoothed density map is overlaid on the corresponding DAPI image for spatial context. Samples are arranged by condition (Control, FSHD1, and DEL5; rows) and by replicate (columns), with all panels shown at the same spatial scale and using the same crop window and colour scale. A 500 $\mu$m scalebar is shown in each panel.

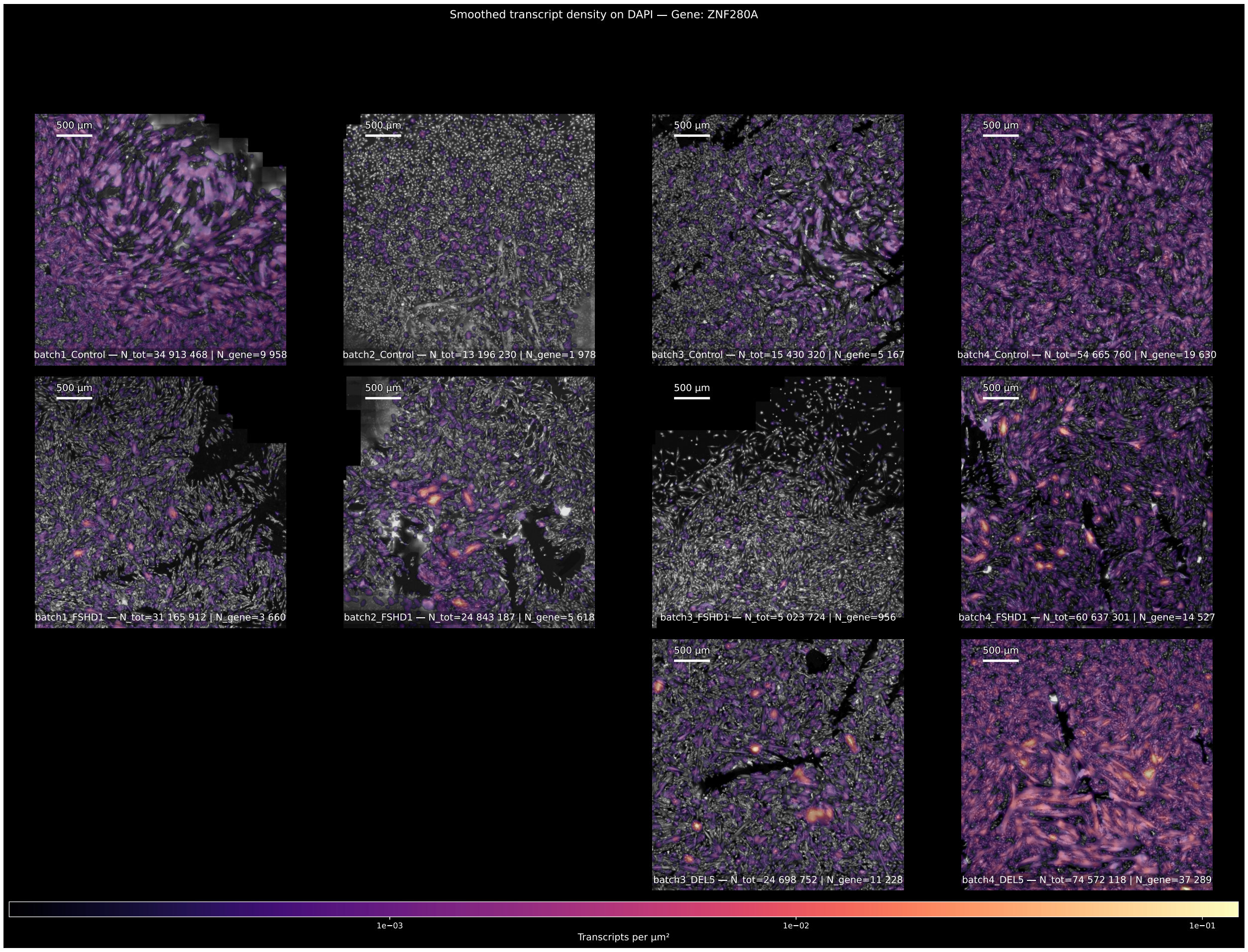


Figure S5: DAPI-normalised spatial density maps for *ZNF280A* in the FSHD MERFISH dataset. For each sample, the raw *ZNF280A* transcript point pattern was transformed into a continuous density field using Gaussian smoothing. The resulting smoothed density map is overlaid on the corresponding DAPI image for spatial context. Samples are arranged by condition (Control, FSHD1, and DEL5; rows) and by replicate (columns), with all panels shown at the same spatial scale and using the same crop window and colour scale. A 500 $\mu$m scalebar is shown in each panel.

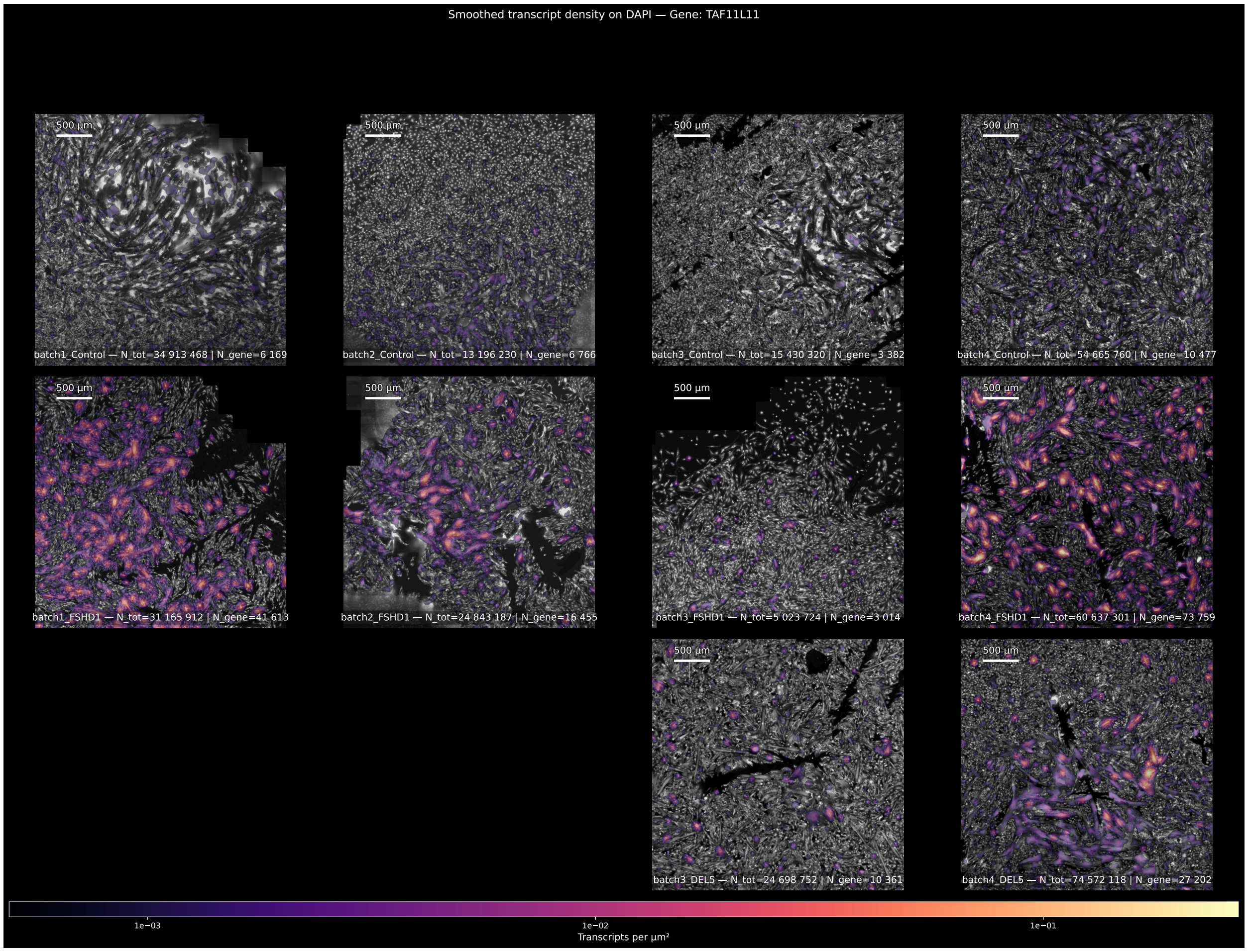


Figure S6: DAPI-normalised spatial density maps for *TAF11L11* in the FSHD MERFISH dataset. For each sample, the raw *TAF11L11* transcript point pattern was transformed into a continuous density field using Gaussian smoothing. The resulting smoothed density map is overlaid on the corresponding DAPI image for spatial context. Samples are arranged by condition (Control, FSHD1, and DEL5; rows) and by replicate (columns), with all panels shown at the same spatial scale and using the same crop window and colour scale. A 500 $\mu$m scalebar is shown in each panel.

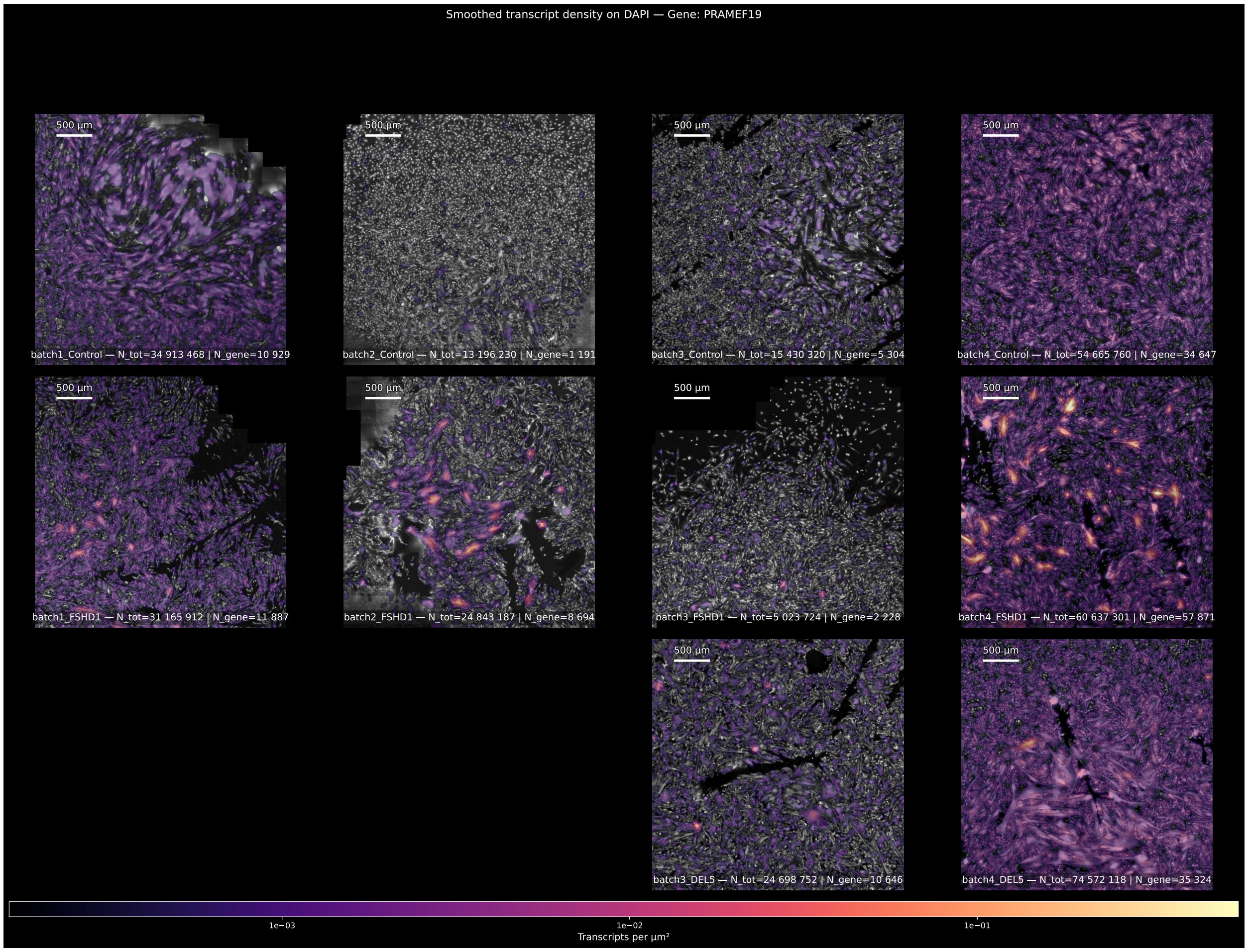


Figure S7: DAPI-normalised spatial density maps for *PRAMEF19* in the FSHD MERFISH dataset. For each sample, the raw *PRAMEF19* transcript point pattern was transformed into a continuous density field using Gaussian smoothing. The resulting smoothed density map is overlaid on the corresponding DAPI image for spatial context. Samples are arranged by condition (Control, FSHD1, and DEL5; rows) and by replicate (columns), with all panels shown at the same spatial scale and using the same crop window and colour scale. A 500 $\mu$m scalebar is shown in each panel.

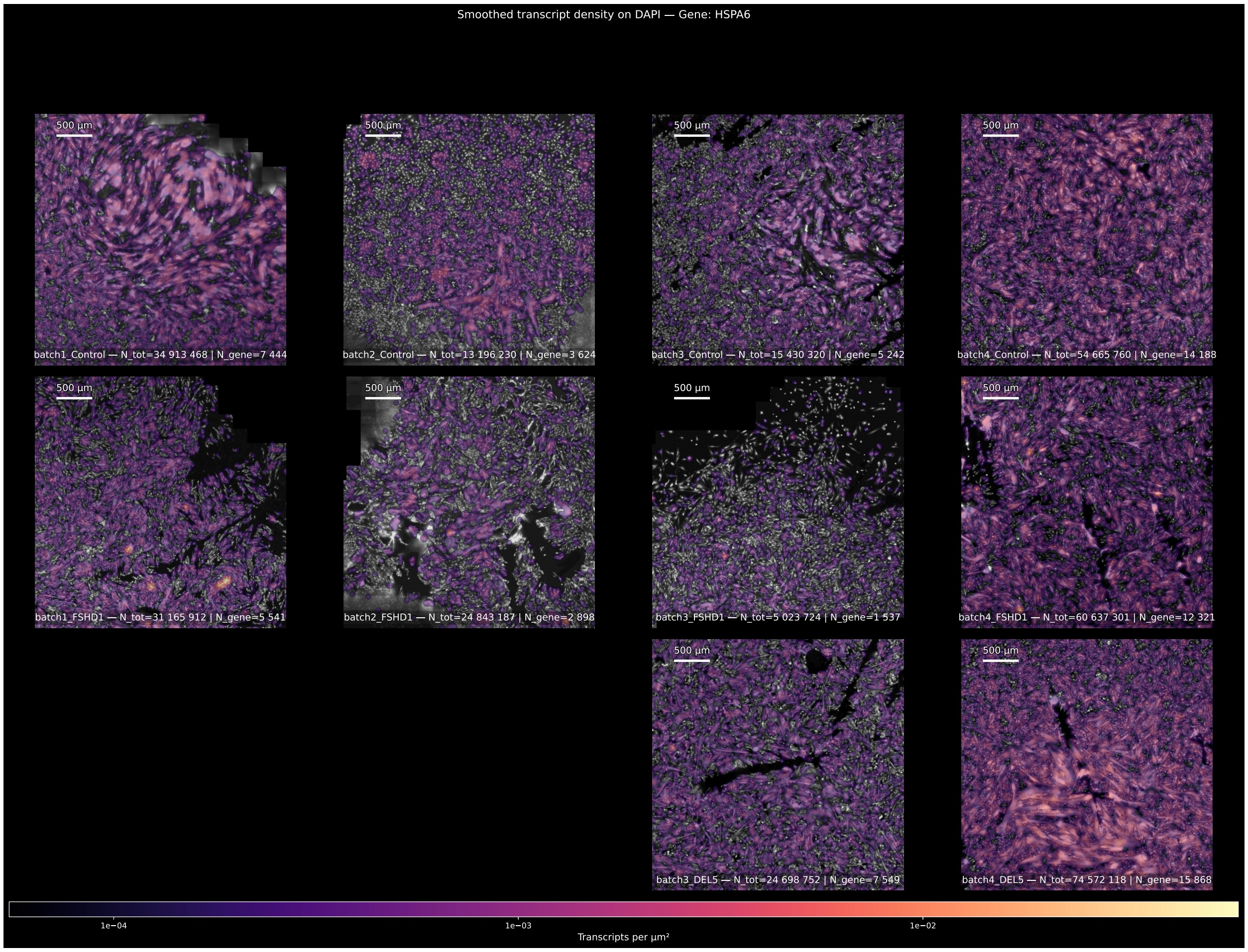


Figure S8: DAPI-normalised spatial density maps for *HSPA6* in the FSHD MERFISH dataset. For each sample, the raw *HSPA6* transcript point pattern was transformed into a continuous density field using Gaussian smoothing. The resulting smoothed density map is overlaid on the corresponding DAPI image for spatial context. Samples are arranged by condition (Control, FSHD1, and DEL5; rows) and by replicate (columns), with all panels shown at the same spatial scale and using the same crop window and colour scale. A 500 $\mu$m scalebar is shown in each panel.

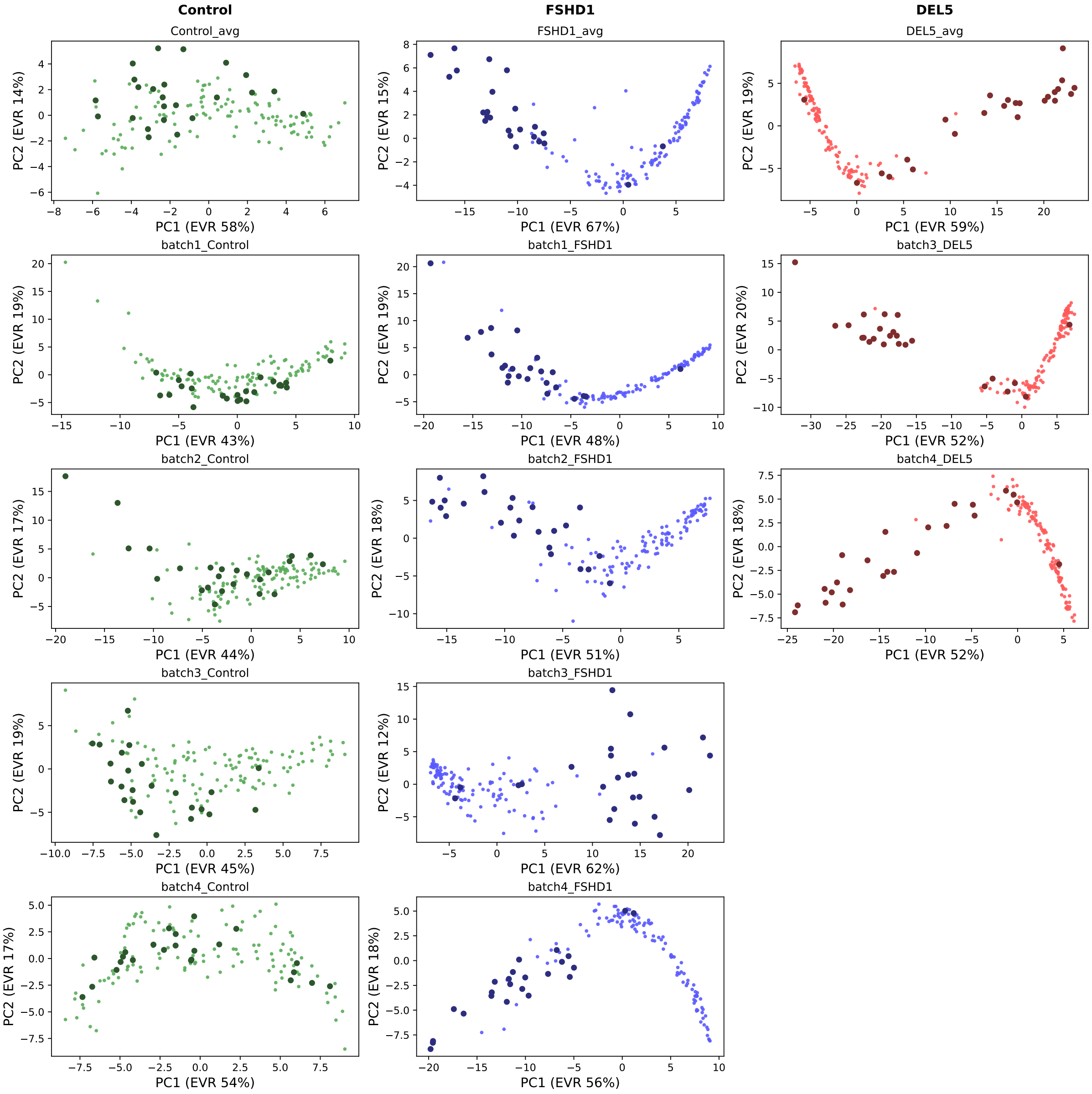


Figure S9: PCA of gene-level Minkowski profiles for the MERFISH FSHD dataset, separated per condition. Each panel shows genes projected onto the first two principal components, computed separately for each individual sample or condition-averaged sample. Columns correspond to conditions (Control, FSHD1, DEL5). The top row shows the condition-averaged samples, and the rows below show individual samples. Each point represents one gene in one averaged condition or in one individual sample. DUX4-target genes are highlighted as darker points within each panel.

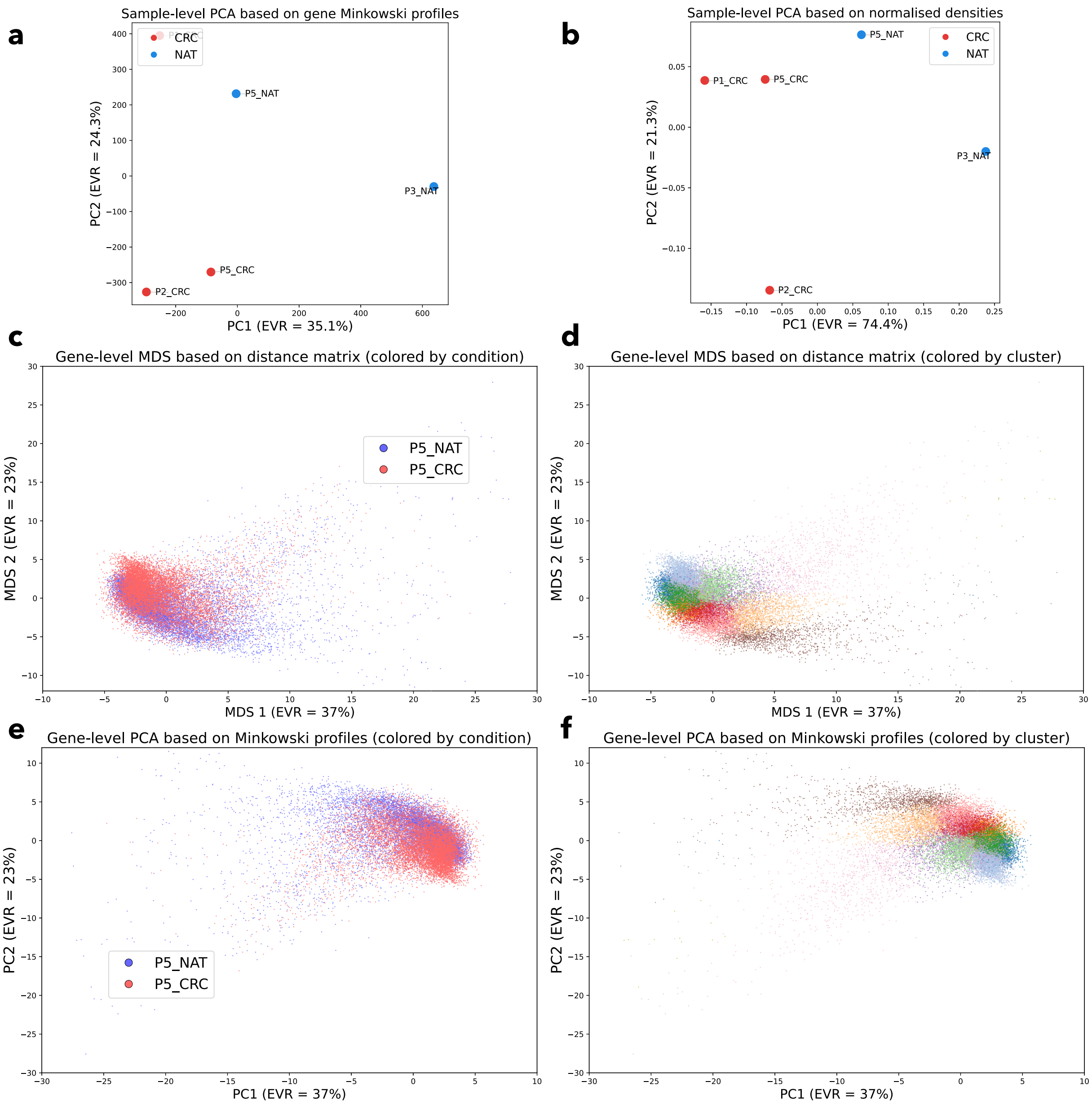


Figure S10: **(a)** PCA of sample-level Minkowski profiles in the CRC Visium HD dataset. Each point represents one sample, coloured by condition. Axis labels indicate the variance explained by PC1 and PC2. **(b)** PCA of sample-level normalised gene densities. Each point represents one sample, projected from the vector of normalised gene densities and coloured by condition. **(c)** Classical MDS embedding of gene-level 2-Wasserstein distances between Minkowski profiles in the P5_CRC and P5_NAT samples. Each point represents one gene in one sample; each gene therefore appears twice, once in P5_CRC and once in P5_NAT. Points are coloured by condition. **(d)** The same MDS embedding, coloured by communities identified with the Leiden algorithm. **(e)** PCA of gene-level Minkowski profiles in the P5_CRC and P5_NAT samples. Each point represents one gene in one sample; each gene therefore appears twice, once in P5_CRC and once in P5_NAT. Points are coloured by condition. Axis labels indicate the variance explained by PC1 and PC2. **(f)** The same PCA embedding, coloured by communities identified with the Leiden algorithm.

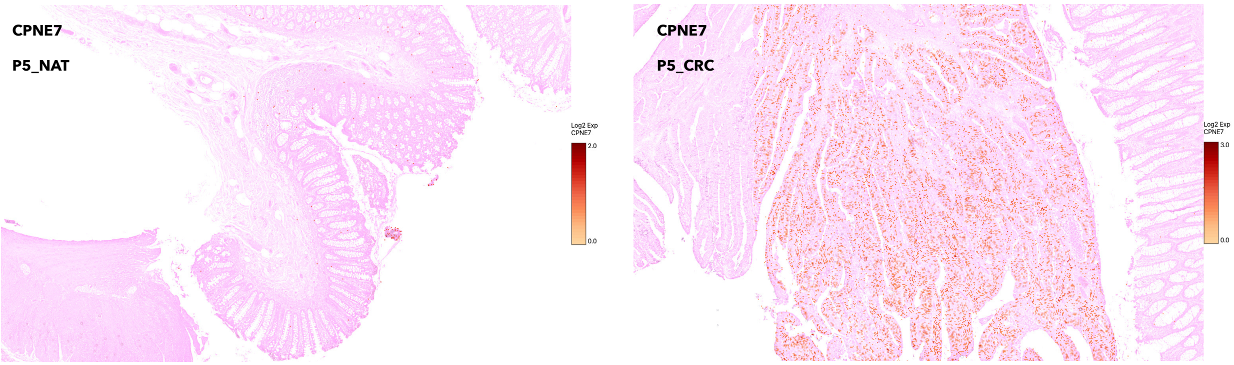


Figure S11: Spatial expression of *CPNE7* [2] visualised with Loupe Browser (10x Genomics) on Visium HD data. Left: P5_NAT sample. Right: P5_CRC sample. The background image corresponds to the histological brightfield tissue image provided with the Visium HD dataset. Transcript abundance for *CPNE7* is overlaid on a grid of 8 $\mu$m pixels, where colour intensity reflects the detected transcript counts per pixel. Both panels are shown on the same colour scale.

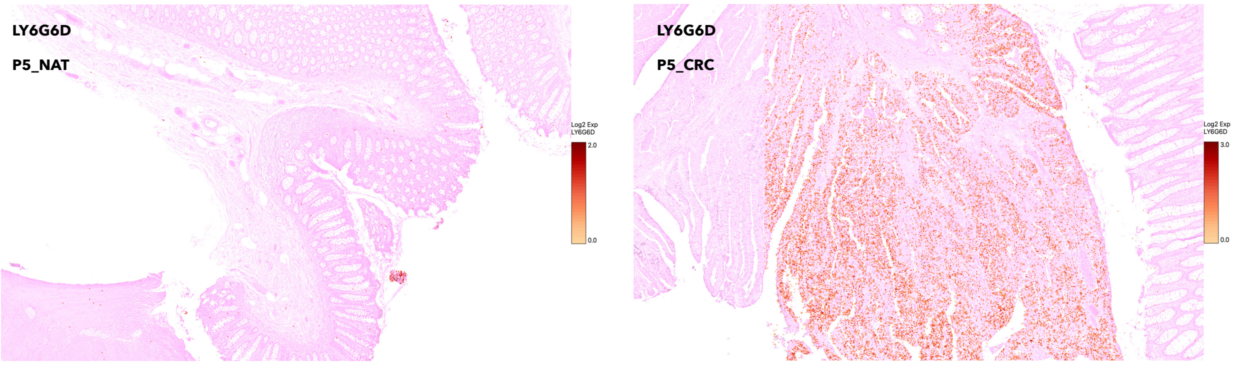


Figure S12: Spatial expression of *LY6G6D* [3] visualised with Loupe Browser (10x Genomics) on Visium HD data. Left: P5_NAT sample. Right: P5_CRC sample. The background image corresponds to the histological brightfield tissue image provided with the Visium HD dataset. Transcript abundance for *LY6G6D* is overlaid on a grid of 8 $\mu$m pixels, where colour intensity reflects the detected transcript counts per pixel. Both panels are shown on the same colour scale.

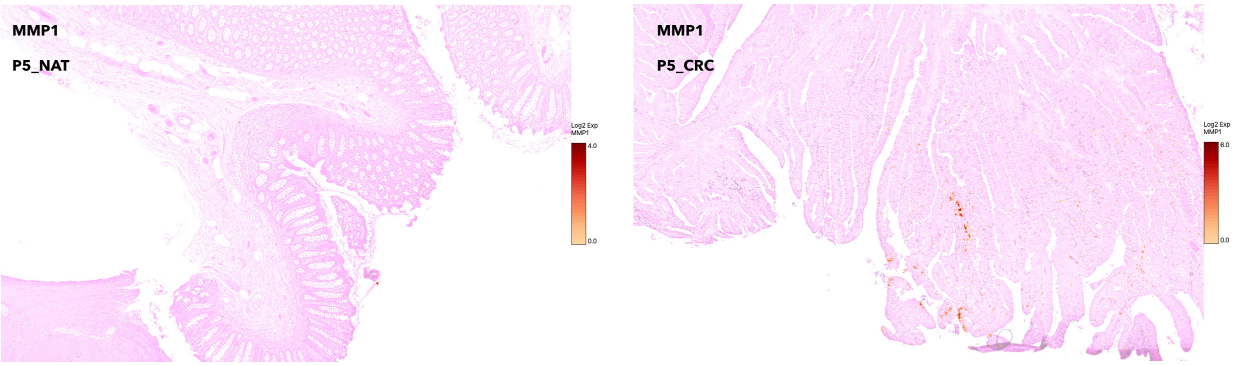


Figure S13: Spatial expression of *MMP1* [4, 5] visualised with Loupe Browser (10x Genomics) on Visium HD data. Left: P5_NAT sample. Right: P5_CRC sample. The background image corresponds to the histological brightfield tissue image provided with the Visium HD dataset. Transcript abundance for *MMP1* is overlaid on a grid of 8 $\mu$m pixels, where colour intensity reflects the detected transcript counts per pixel. Both panels are shown on the same colour scale.

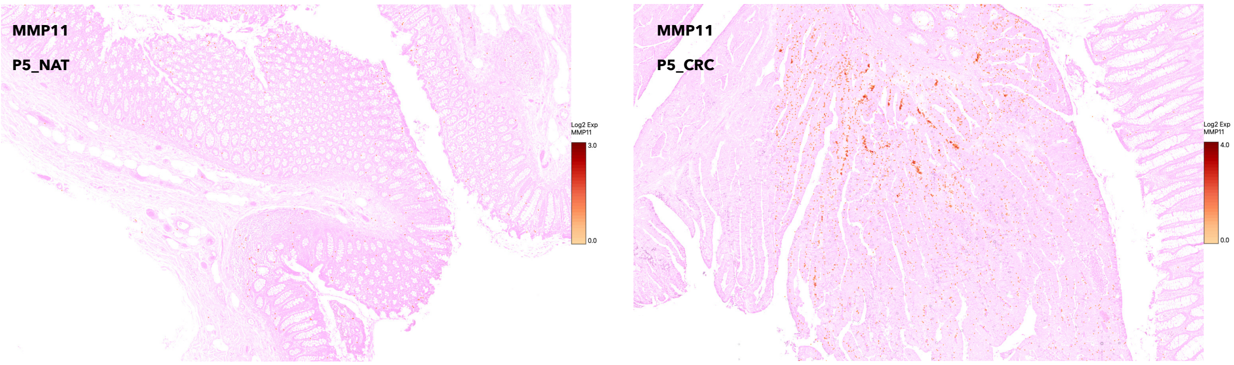


Figure S14: Spatial expression of *MMP11* [4, 5] visualised with Loupe Browser (10x Genomics) on Visium HD data. Left: P5_NAT sample. Right: P5_CRC sample. The background image corresponds to the histological brightfield tissue image provided with the Visium HD dataset. Transcript abundance for *MMP11* is overlaid on a grid of 8 $\mu$m pixels, where colour intensity reflects the detected transcript counts per pixel. Both panels are shown on the same colour scale.

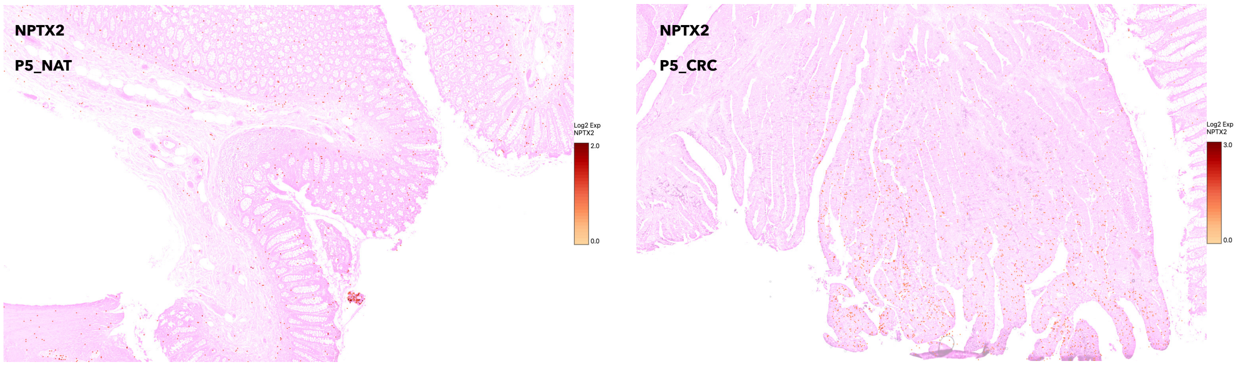


Figure S15: Spatial expression of *NPTX2* [6] visualised with Loupe Browser (10x Genomics) on Visium HD data. Left: P5_NAT sample. Right: P5_CRC sample. The background image corresponds to the histological brightfield tissue image provided with the Visium HD dataset. Transcript abundance for *NPTX2* is overlaid on a grid of 8 $\mu$m pixels, where colour intensity reflects the detected transcript counts per pixel. Both panels are shown on the same colour scale.

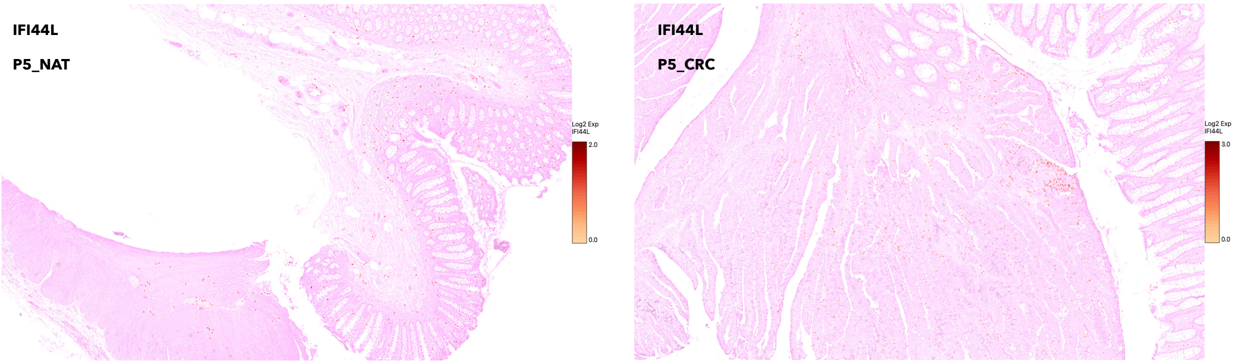


Figure S16: Spatial expression of *IFI44L* [7] visualised with Loupe Browser (10x Genomics) on Visium HD data. Left: P5_NAT sample. Right: P5_CRC sample. The background image corresponds to the histological brightfield tissue image provided with the Visium HD dataset. Transcript abundance for *IFI44L* is overlaid on a grid of 8 $\mu$m pixels, where colour intensity reflects the detected transcript counts per pixel. Both panels are shown on the same colour scale.

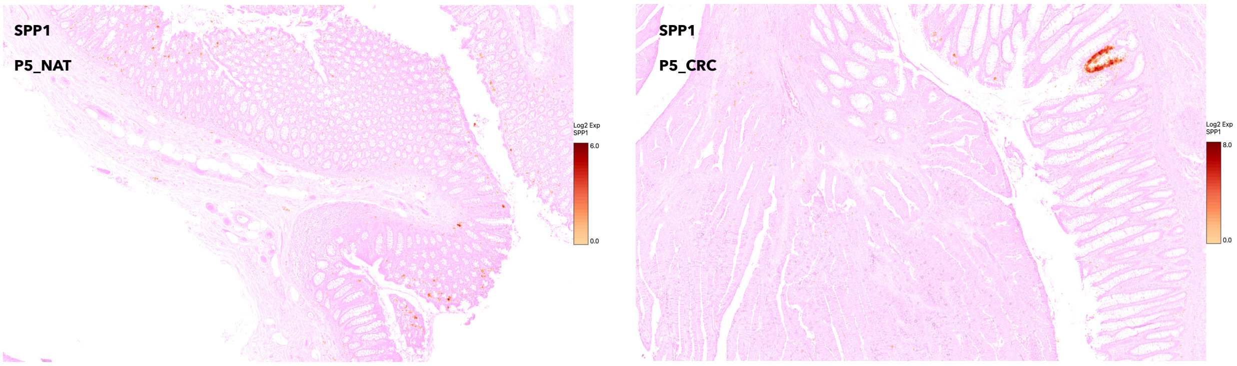


Figure S17: Spatial expression of *SPP1* visualised with Loupe Browser (10x Genomics) on Visium HD data. Left: P5_NAT sample. Right: P5_CRC sample. The background image corresponds to the histological brightfield tissue image provided with the Visium HD dataset. Transcript abundance for *SPP1* is overlaid on a grid of 8 $\mu$m pixels, where colour intensity reflects the detected transcript counts per pixel. Both panels are shown on the same colour scale.

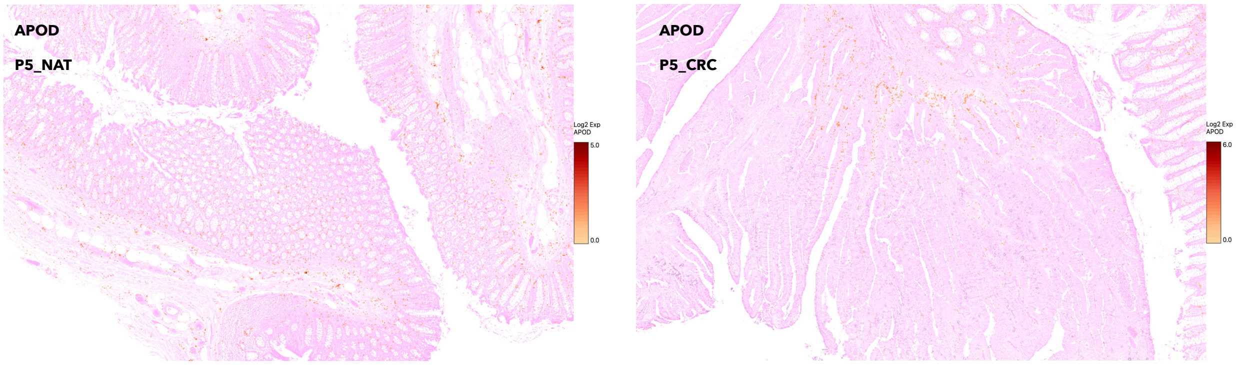


Figure S18: Spatial expression of *APOD* visualised with Loupe Browser (10x Genomics) on Visium HD data. Left: P5_NAT sample. Right: P5_CRC sample. The background image corresponds to the histological brightfield tissue image provided with the Visium HD dataset. Transcript abundance for *APOD* is overlaid on a grid of 8 $\mu$m pixels, where colour intensity reflects the detected transcript counts per pixel. Both panels are shown on the same colour scale.

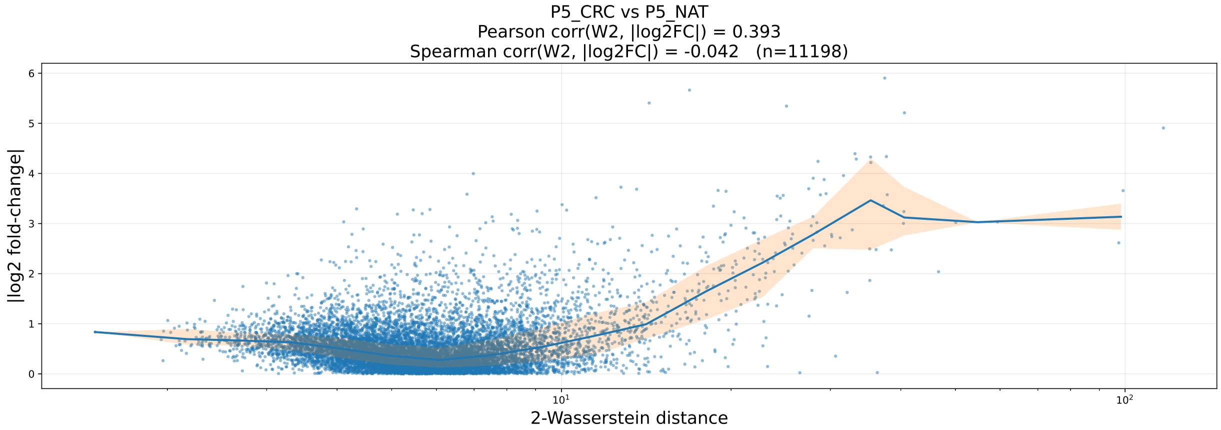


Figure S19: Relationship between spatial reorganisation and differential gene expression in the CRC Visium HD dataset. The panel shows the gene-level 2-Wasserstein distance between Minkowski profiles plotted against the absolute $\log_2$ fold change computed from normalised gene densities. Each point represents one gene. The solid curve represents the median $|\log_2 \mathrm{FC}|$ within logarithmic bins of 2-Wasserstein distances, and the shaded region indicates the interquartile range. The Pearson and Spearman correlations between 2-Wasserstein distances and $|\log_2 \mathrm{FC}|$ are reported in the panel.

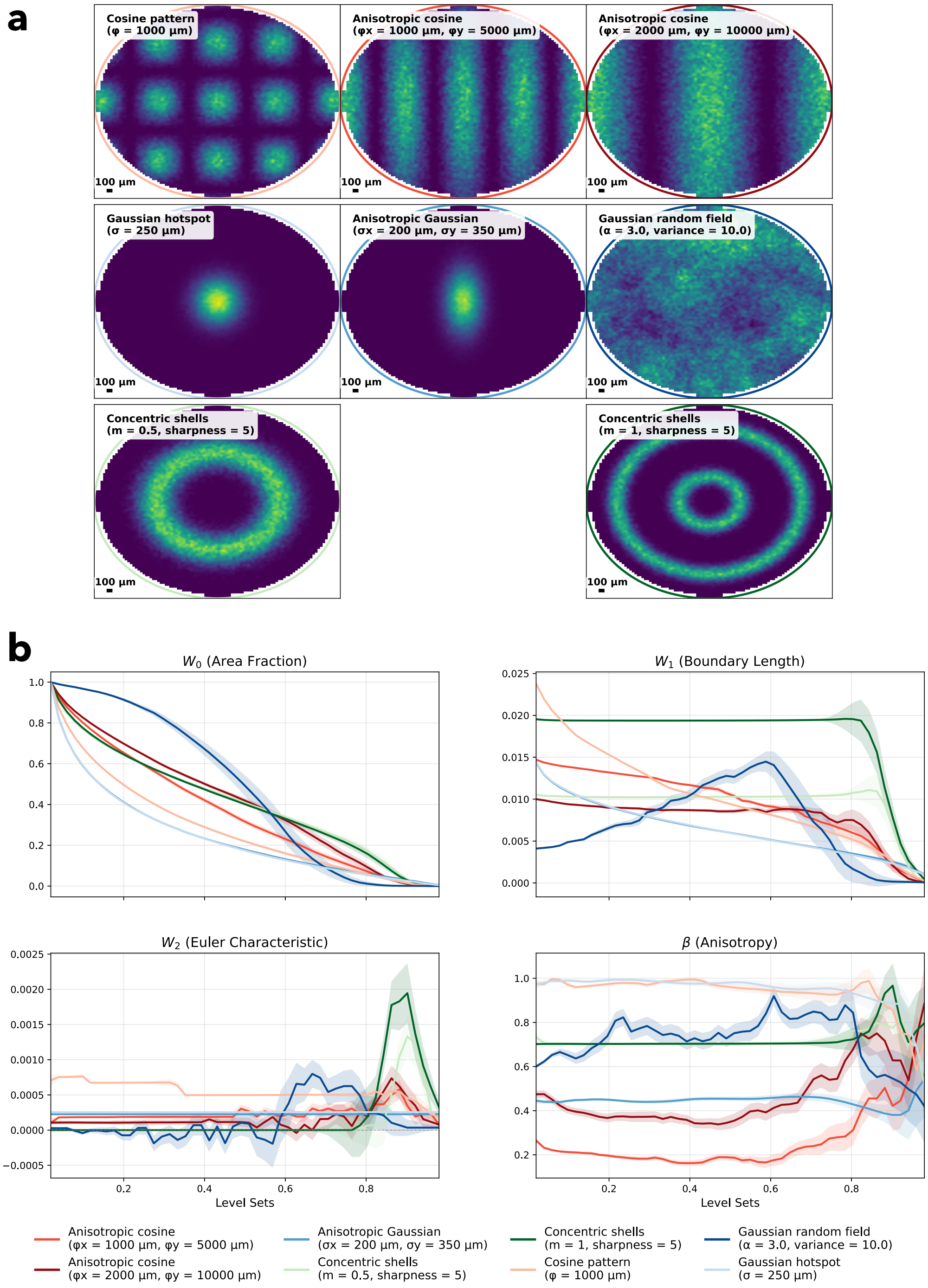


Figure S20: **(a)** Synthetic point distributions used to illustrate the Minkowski characteristics. Eight toy models are generated within a common elliptical mask and arranged by family: cosine-based patterns (top row), Gaussian-based patterns (middle row), and concentric-shell patterns (bottom row). Each panel shows the smoothed transcript density as a colourmap. The ellipse boundary is colour-coded by family, using distinct shades of red (cosines), blue (Gaussians, including the Gaussian random field), and green (concentric shells). **(b)** Corresponding estimated Minkowski characteristics across density level sets for the same eight toy models. Solid lines indicate the estimated Minkowski characteristics; shaded bands show ±1 standard deviation estimated from Monte Carlo resampling.

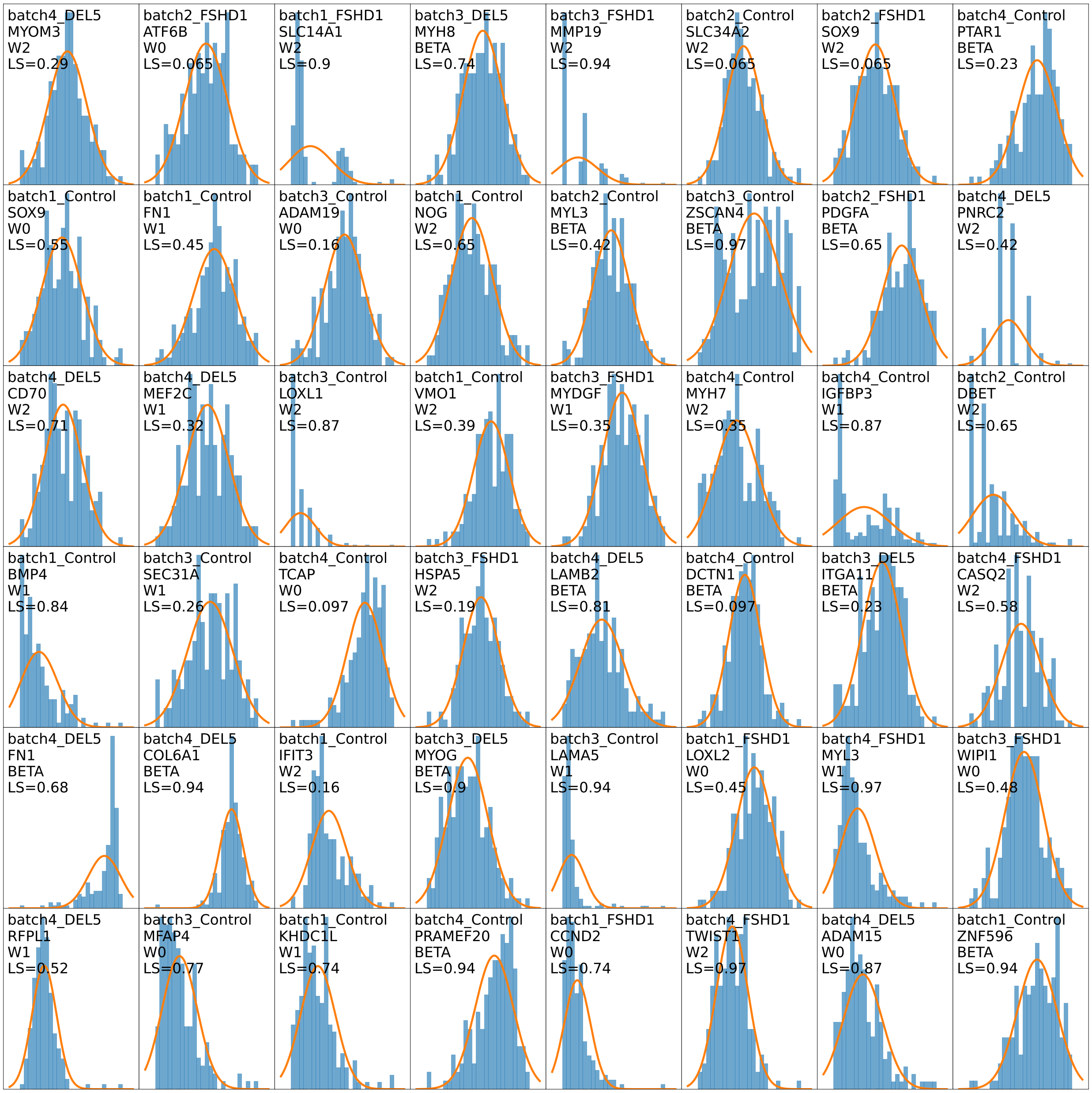


Figure S21: Visual normality diagnostic for Monte Carlo resamples of Minkowski profile components. Mosaic of randomly selected tuples (sample, gene, Minkowski characteristic, level set): each panel displays the empirical distribution of the corresponding Monte Carlo resamples as a density-normalised histogram. The orange curve shows the moment-matched Gaussian density (same mean and variance as the Monte Carlo samples). This qualitative check supports the use of a closed-form Gaussian 2-Wasserstein distance.

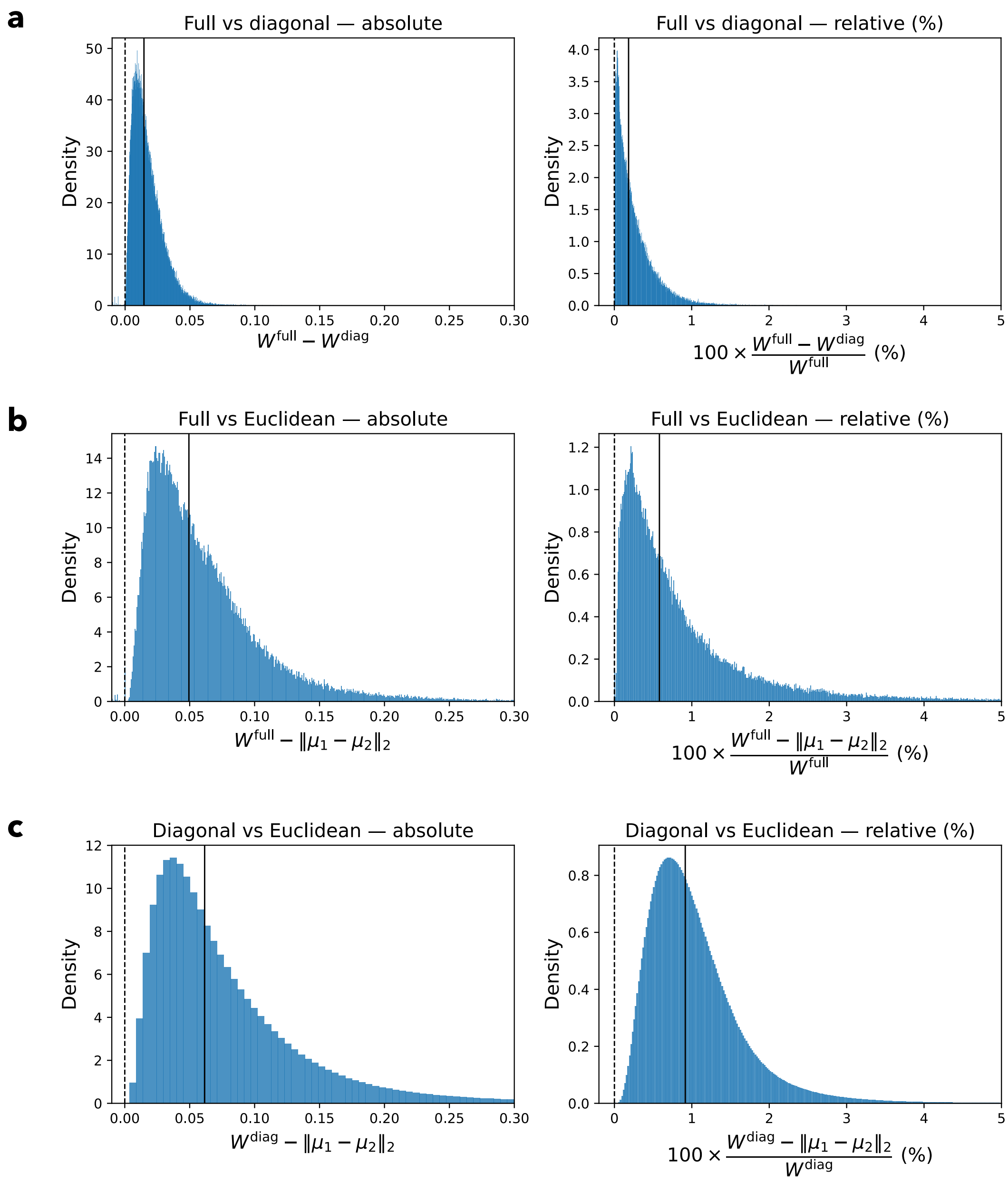


Figure S22: Distribution of absolute (left) and relative (right) differences between distances computed from Minkowski profiles. **(a,b)** Differences aggregated over all combinations of gene pairs in the DEL5, FSHD1 and Control averaged samples of the MERFISH FSHD dataset. **(a)** Full-covariance Gaussian 2-Wasserstein distance between Minkowski profiles compared with its diagonal approximation, which ignores cross covariance terms. **(b)** Full-covariance Gaussian 2-Wasserstein distance compared with the Euclidean distance between Minkowski profiles. **(c)** Differences aggregated over all gene pairs in the P5_CRC and P5_NAT samples of the Visium HD colorectal cancer dataset, comparing the diagonal Gaussian 2-Wasserstein distance with the Euclidean distance between Minkowski profiles. Relative differences are reported as $100 \times (W^{\text{ref}} - W^{\text{approx}})/W^{\text{ref}}$, where $W^{\text{ref}} = W^{\text{full}}$ in panels **(a,b)** and $W^{\text{ref}} = W^{\text{diag}}$ in panel **(c)**. Solid black lines indicate the medians of the displayed distributions, and dashed black lines indicate zero difference.

#### S4 Supplementary Tables

Table S1: Community summary for the FSHD MERFISH dataset. For each community, we report the number of nodes and unique genes, the retention (percentage of genes assigned to the same community across all averaged conditions), the proportion of DUX4-target (GOI: Genes Of Interest) nodes and the proportion of DUX4-target genes covered, as well as the community composition by condition (node percentages) and the mean normalised gene density within each condition.

| Community | $n_{\text{nodes}}$ | $n_{\text{genes}}$ | Retention [%] | GOI nodes [%] | GOI genes [%] | Control_avg [%] | Control mean density | FSHD1_avg [%] | FSHD1 mean density | DEL5_avg [%] | DEL5 mean density |
|---|---|---|---|---|---|---|---|---|---|---|---|
| 0 | 204 | 105 | 31.4 | 11.3 | 91.7 | 35.3 | $9.17 \times 10^{-3}$ | 30.9 | $6.09 \times 10^{-3}$ | 33.8 | $1.21 \times 10^{-2}$ |
| 1 | 162 | 79 | 40.5 | 4.32 | 25.0 | 42.0 | $9.35 \times 10^{-3}$ | 27.8 | $8.54 \times 10^{-3}$ | 30.2 | $1.40 \times 10^{-2}$ |
| 2 | 54 | 33 | 0.000 | 77.8 | 91.7 | 0.000 | $1.33 \times 10^{-3}$ | 59.3 | $4.63 \times 10^{-3}$ | 40.7 | $2.99 \times 10^{-3}$ |

Table S2: Community summary for the CRC Visium HD dataset. For each community, we report the number of nodes and unique genes, the retention (percentage of genes assigned to the same community across both conditions), the community composition by condition (node percentages), and the mean normalised gene density within each condition.

| Community | $n_{\text{nodes}}$ | $n_{\text{genes}}$ | Retention [%] | P5_NAT [%] | P5_NAT mean density | P5_CRC [%] | P5_CRC mean density |
|---|---|---|---|---|---|---|---|
| 0 | 2261 | 2128 | 6.25 | 85.45 | $1.01 \times 10^{-3}$ | 14.55 | $7.39 \times 10^{-4}$ |
| 1 | 2067 | 2016 | 2.53 | 3.68 | $1.08 \times 10^{-3}$ | 96.32 | $5.54 \times 10^{-4}$ |
| 2 | 1964 | 1863 | 5.42 | 85.54 | $3.45 \times 10^{-4}$ | 14.46 | $2.40 \times 10^{-4}$ |
| 3 | 1888 | 1657 | 13.94 | 48.73 | $1.29 \times 10^{-4}$ | 51.27 | $8.45 \times 10^{-5}$ |
| 4 | 1758 | 1736 | 1.27 | 5.97 | $5.62 \times 10^{-4}$ | 94.03 | $4.21 \times 10^{-4}$ |
| 5 | 1692 | 1658 | 2.05 | 9.93 | $2.23 \times 10^{-4}$ | 90.07 | $1.38 \times 10^{-4}$ |
| 6 | 1636 | 1534 | 6.65 | 47.25 | $4.38 \times 10^{-4}$ | 52.75 | $3.63 \times 10^{-4}$ |
| 7 | 1621 | 1421 | 14.07 | 73.84 | $1.71 \times 10^{-4}$ | 26.16 | $1.21 \times 10^{-4}$ |
| 8 | 1485 | 1322 | 12.33 | 59.26 | $1.32 \times 10^{-3}$ | 40.74 | $7.96 \times 10^{-4}$ |
| 9 | 1449 | 1432 | 1.19 | 7.80 | $1.44 \times 10^{-3}$ | 92.20 | $8.96 \times 10^{-4}$ |
| 10 | 1344 | 1191 | 12.85 | 81.55 | $1.05 \times 10^{-4}$ | 18.45 | $6.30 \times 10^{-5}$ |
| 11 | 1199 | 1152 | 4.08 | 81.90 | $8.66 \times 10^{-4}$ | 18.10 | $6.33 \times 10^{-4}$ |
| 12 | 1004 | 967 | 3.83 | 67.73 | $2.71 \times 10^{-4}$ | 32.27 | $1.82 \times 10^{-4}$ |
| 13 | 941 | 722 | 30.33 | 54.41 | $2.81 \times 10^{-4}$ | 45.59 | $1.06 \times 10^{-4}$ |
| 14 | 54 | 50 | 8.00 | 85.19 | $1.69 \times 10^{-4}$ | 14.81 | $3.25 \times 10^{-5}$ |
| 15 | 24 | 24 | 0.00 | 100.00 | $3.46 \times 10^{-4}$ | 0.00 | $3.16 \times 10^{-5}$ |
| 16 | 12 | 12 | 0.00 | 100.00 | $1.49 \times 10^{-4}$ | 0.00 | $9.38 \times 10^{-6}$ |
| 17 | 3 | 3 | 0.00 | 66.67 | $6.06 \times 10^{-4}$ | 33.33 | $5.63 \times 10^{-5}$ |
| 18 | 3 | 3 | 0.00 | 100.00 | $2.58 \times 10^{-4}$ | 0.00 | $8.38 \times 10^{-6}$ |
| 19 | 1 | 1 | 0.00 | 100.00 | $2.68 \times 10^{-4}$ | 0.00 | $1.70 \times 10^{-5}$ |
| 20 | 1 | 1 | 0.00 | 100.00 | $1.48 \times 10^{-3}$ | 0.00 | $3.16 \times 10^{-6}$ |
| 21 | 1 | 1 | 0.00 | 100.00 | $1.21 \times 10^{-3}$ | 0.00 | $6.77 \times 10^{-4}$ |

### References


[1] Pang-Ning Tan, Michael Steinbach, Anuj Karpatne, and Vipin Kumar. *Introduction to Data Mining.* Pearson, Boston, 2 edition, 2018.

[2] Liangbo Zhao, Xiao Sun, Chenying Hou, Yanmei Yang, Peiwen Wang, Zhaoyuan Xu, Zhenzhen Chen, Xiangrui Zhang, Guanghua Wu, Hong Chen, et al. Cpne7 promotes colorectal tumorigenesis by interacting with nono to initiate zfp42 transcription. *Cell Death & Disease*, 15(12):896, 2024.

[3] Guido Giordano, Pietro Parcesepe, Mario Rosario D'Andrea, Luigi Coppola, Tania Di Raimo, Andrea Remo, Erminia Manfrin, Claudia Fiorini, Aldo Scarpa, Carla Azzurra Amoreo, et al. Jak/stat5-mediated subtype-specific lymphocyte antigen 6 complex, locus g6d (ly6g6d) expression drives mismatch repair proficient colorectal cancer. *Journal of Experimental & Clinical Cancer Research*, 38(1):28, 2019.

[4] Graeme I Murray, Margaret E Duncan, Pauline O'Neil, William T Melvin, and John E Fothergill. Matrix metalloproteinase–1 is associated with poor prognosis in colorectal cancer. *Nature medicine*, 2(4):461–462, 1996.

[5] Jing Yu, Zhen He, Xiaowen He, Zhanhao Luo, Lei Lian, Baixing Wu, Ping Lan, and Haitao Chen. Comprehensive analysis of the expression and prognosis for mmps in human colorectal cancer. *Frontiers in Oncology*, 11:771099, 2021.

[6] Chunjie Xu, Guangang Tian, Chunhui Jiang, Hanbing Xue, Manzila Kuerbanjiang, Longci Sun, Lei Gu, Hong Zhou, Ye Liu, Zhigang Zhang, et al. Nptx2 promotes colorectal cancer growth and liver metastasis by the activation of the canonical wnt/$\beta$-catenin pathway via fzd6. *Cell death & disease*, 10(3):217, 2019.

[7] John W Schoggins, Sam J Wilson, Maryline Panis, Mary Y Murphy, Christopher T Jones, Paul Bieniasz, and Charles M Rice. A diverse range of gene products are effectors of the type i interferon antiviral response. *Nature*, 472(7344):481–485, 2011.


$LN@col2$
